# *APOE* ɛ2 vs *APOE* ɛ4 dosage shows sex-specific links to hippocampus-default network subregion co-variation

**DOI:** 10.1101/2022.03.15.484482

**Authors:** Chloé Savignac, Sylvia Villeneuve, AmanPreet Badhwar, Karin Saltoun, Kimia Shafighi, Chris Zajner, Vaibhav Sharma, Sarah A Gagliano Taliun, Sali Farhan, Judes Poirier, Danilo Bzdok

## Abstract

Alzheimer’s disease and related dementias (ADRD) are marked by intracellular tau aggregates in the medial-temporal lobe (MTL) and extracellular amyloid aggregates in the default network (DN). Here, we sought to clarify ADRD-related co-dependencies between the MTL’s most vulnerable structure, the hippocampus (HC), and the highly associative DN at a subregion resolution. We confronted the effects of *APOE* ɛ2 and ɛ4, rarely investigated together, with their impact on HC-DN co-variation regimes at the population level. In a two-pronged decomposition of structural brain scans from ∼40,000 UK Biobank participants, we located co-deviating structural patterns in HC and DN subregions as a function of ADRD family risk. Across the disclosed HC-DN signatures, recurrent deviations in the CA1, CA2/3, molecular layer, fornix’s fimbria, and their cortical partners related to ADRD risk. Phenome-wide profiling of HC-DN co- variation expressions from these population signatures revealed male-specific associations with air-pollution, and female-specific associations with cardiovascular traits. We highlighted three main factors associated with brain-*APOE* associations across the different gene variants: happiness, and satisfaction with friendships, and with family. We further showed that *APOE* ɛ2/2 interacts preferentially with HC-DN co-variation patterns in estimating social lifestyle in males and physical activity in females. Our findings reinvigorate the often-neglected interplay between *APOE* ɛ2 dosage and sex, which we have linked to fine-grained structural divergences indicative of ADRD susceptibility.

## Introduction

Around the globe, >50 million people are living with dementia – a global burden of >1 trillion USD$ annually [1]. By 2050, an estimated threefold increase in affected individuals is projected as a result of increased longevity [2]. The anticipated explosion in the number of dementia cases will put a strain on the 82 billion hours of annual informal care provided by caretakers worldwide [1]. In contrast to this secular trend, the age-specific prevalence of dementia is expected to decrease in certain high-income countries, which can be attributable to improvement in underlying health and socioeconomic determinants [2]. A recent authoritative report on dementia prevention has identified about a dozen potentially modifiable risk factors that could explain disparity in ADRD incidence [3]. The disparate risk dimensions include personal habits and lifestyle, physical and mental health, as well as societal and external factors. New public health policies targeted at reducing mid- to late-life risk factors (e.g., physical inactivity, social disengagement, loneliness) thus have the potential to delay dementia onset in the most deprived older adults. As the global prevalence of dementia is quickly rising, there is an unpreceded need to characterize the impact of genetic predisposition (e.g., Apolipoprotein E (*APOE*) polymorphism [4]) and modifiable risk factors on ADRD-vulnerable brain structures before the onset of cognitive downwards drifts.

Over the past two decades, brain-imaging studies have converged on disruption of a coherent network of higher association regions that involve key nodes of the default network (DN) in individuals with ADRD compared to healthy controls [5]. Extensive efforts have mobilized resting-state functional connectivity analyses to investigate patients with ADRD, with converging results in the DN [6]. However, delineating a definitive profile of functional connectivity deviations related to ADRD risk in healthy subjects was plagued with slow progress. Most such biomarker studies have attempted to identify functional connectivity patterns that reliably tell apart ɛ4 carriers from non-carriers. Yet, most other *APOE* variants have been largely neglected, perhaps because they occur much more infrequently in the general population. The extensive literature on altered DN connectivity in ɛ4 carriers has yet to reach consensus as reports of both increased [7] and decreased [8] connectivity within nodes of the DN have repeatedly led to contradictory conclusions. Among the few studies that could investigate concurrent connectivity alterations in the hippocampus (HC) and regions of the DN in ɛ4 carriers, the HC was typically treated as a monolithic structure [9], rather than appreciating its functional and structural heterogeneity. That is, it was studied as a single node when interrogating its coupling links to other DN nodes [10]. These inconsistencies are probably also due in part to data acquisition and preprocessing methods for functional connectivity analysis, which have made some findings in ɛ4 carriers hard to replicate [11]. Moreover, because of the overwhelming singular focus on ɛ4 carriers in the research community, the neural correlates associated with other *APOE* variants remain underspecified. Of particular appeal, illuminating the allegedly opposing effects of *APOE* ɛ2 and ɛ4 on DN and HC integrity could be crucial in guiding potential treatment avenues, given the ɛ2-associated protective outcome on brain structure [12].

A parallel stream of literature has focused on changes in hippocampal microstructure over the course of ADRD progression, mostly by performing thorough post-mortem autopsy on patients with probable ADRD. The hippocampus formation is known for subfield-specific vulnerability to ADRD, at least since the late 1990s [13]. Yet, the hippocampus is still routinely treated as if it was an anatomically homogeneous structure in common brain-imaging studies [9, 14, 15]. By extension, such an analytical approach is blind to the distinct links between HC subregions and DN subregions. For example, the presubiculum and parasubiculum receive important axon projections from the retrosplenial cortex and posterior cingulate cortex [16]. Yet, the fornix, which carries the axons from the CA1 and subiculum, forwards the only hippocampal output signals that directly go to the ventromedial and orbitofrontal cortex of the DN [17, 18]. Glossing over these known microanatomical nuances could explain reports of poor predictive value of hippocampal atrophy in early ADRD stages when measuring analyzing the whole hippocampus as a single unit. In a randomized clinical trial, baseline hippocampus volumes, manually traced and corrected for inhomogeneity, predicted conversion to ADRD over a 3-year period at 60.4% accuracy [19]. With the advent of ultra-high-resolution atlases and advanced automatic sub-segmentation techniques, assessment of the subfield-specific vulnerably of both hippocampi to ADRD progression in an observer-independent fashion is now coming into reach. Instead of relying mostly on post-mortem autopsy from patients to ultimately confirm ADRD status, we will soon be able to directly, non-invasively, quantify the level of risk of a given patient based on subfield-level granular information. From the perspective of clinical translation, coming up with individual profiles of microstructural alterations that are characteristic of ADRD risk could usher towards a principled path towards precision medicine in neurology.

For these reasons, we here opted for structural brain-imaging to relate genetic risk to robust co-dependence principles between neocortical DN and allocortical HC at subregion granularity. Given the panoply of individual factors that may affect cortical blood flow (e.g., vigilance, mood, cortisol levels, and coffee intake), functional connectivity would be likely to paint a more circumstantial portrait of ADRD vulnerability. We therefore designed an analytical framework for two-pronged decomposition to zoom in on the structural correspondence between HC and DN subregions at the population level. The doubly multivariate approach was carefully tailored to derive coherent signatures of HC-DN co-variation that are sensitive to the subregion-specific vulnerability of these neural circuits in ADRD. We were able to quantify the level of risk by looking for structural deviation in individuals with and without family history of ADRD by deep inspection of concomitant regimes of HC-DN co-variation. Capitalizing on the rich phenotyping available for 40,000 UK Biobank participants, our study could confront the effects of *APOE* ɛ2 and ɛ4 on inter-individual expressions of HC-DN co-variation — something out of reach for in traditional brain-imaging studies in small to medium sample sizes. In so doing, our study was also uniquely positioned to illuminate possible sex-specific associations across less prevalent *APOE* gene variants that previous brain-imaging investigations systematically ignored.

## Results

### Rationale

In post-mortem autopsy of patients with ADRD, structural alterations of microanatomically defined subregions composing the human HC have been described *in extenso* [20]. Despite such insights from rigorous invasive studies, the overwhelming majority of existing brain-imaging studies has treated the HC as a monolithic brain structure. Hence, the specific vulnerability of its heterogeneous subregions to ADRD pathology remains largely concealed today. Advances in automatic segmentation techniques for the HC using ex vivo brain-imaging allow for subject-specific parcellations that respect the diversity of distinct subregions identified post-mortem. Capitalizing on these ultra-high resolution segmentations, we are now equipped to assess microstructural alteration of the human HC in a newly detailed way that scales to the ∼40,000 UKB participants [21]. These advances enabled us to describe ADRD-related patterns of structural co-variation in DN subregions which were in lockstep with fine-grained HC subregions. Working at population scale made it possible for us to investigate the effect of rare genotypes on brain structure. This approach was especially fruitful for the less common *APOE* ɛ2/2, which has a prevalence of <1% amongst the general population [22]. Given this setup, our investigation was uniquely positioned to carry out sex-specific examinations across all *APOE* gene variants that previous brain-imaging studies systematically ignored. The availability of deep profiling of the UKB participants further allowed us to chart brain-behaviour associations across the whole phenome in an impartial data-driven approach.

### Population signatures of HC-DN co-variation capture subregion-level structural ties

We first delineated the structural dependencies between the subregion atlas of the HC and that of the DN to identify deviations that jointly go hand-in-hand. We benefitted from CCA, a doubly multivariate pattern-learning tool (cf. methods), to identify the sources of common population variation between the full sets of 38 HC subregions and that of 91 DN subregions. This algorithmic approach finds principled signatures of structural co-variation between two sets of variables [23]. Patterns of shared co-variation (*canonical variates,* cf. methods) embed the effects of HC or DN subregion sets in a new representational space where the two sets were most strongly correlated with each other. Pairs of canonical variates, one for the HC and one for the DN, are what we henceforth call *modes.* By construction, these are ranked by importance; and each mode carries unique information by being uncorrelated from each other. Each mode thus represented a different brain *signature* that accounted for increasingly less shared variance between the neocortical and allocortical atlas at subregion resolution.

We focused on the leading 25 modes, mode 1 being the most explanatory signature of HC-DN co-variation under the elected model. The explanatory power by a given mode was quantified by Pearson’s correlation between inter-individual variation tracked by its associated HC and DN patterns (*canonical correlation*, cf. method). The leading signature of HC-DN co- variation (mode 1) achieved a canonical correlation of rho = 0.51, whereas the second and third signatures achieved correlations of rho = 0.42 and 0.39, respectively. Canonical correlations accounted for increasingly less joint variation between the HC and DN subregions up to the last signature (mode 25), which achieved a correlation of rho = 0.06. The full list of correlation coefficients for the remaining modes has been published elsewhere [24] and is openly accessible online (https://figshare.com/articles/figure/Loneliness_Suppplement_July_22_docx/15060684). This multivariate decomposition served as the backbone for all subsequent analyses that aimed to elucidate how individual expressions of HC-DN co-variation varied in relation to ADRD risk.

### Signatures of HC-DN co-variation illuminate concomitant deviations in ADRD risk

To interrogate the neurobiological manifestations of ADRD family history in our UKB cohort, we performed a rigorous group difference analysis that highlighted any statistically robust ADRD-related divergences in each HC-DN population signature. In doing so, we uncovered the precise subset of anatomical subregions contributing to structural HC-DN co-variation that systematically diverged in individuals with vs without family history of ADRD. A HC or DN subregion observed to have a robustly different co-variation expression in individuals with and without family history of ADRD is henceforth termed a *hit*. We observed a total of 28 HC and 135 DN hits across the leading 25 modes. As a general trend, HC hits were mainly located in the cornu ammonis (CA) subregions (42.9% of total divergences). Parallel DN hits were predominantly observed in the prefrontal cortex (dorsomedial prefrontal cortex (dmPFC) and ventrolateral prefrontal cortex (vPFC); 45.9% of total divergences), and posterior midlines structures (posterior cingulate cortex (PCC), precuneus (PCu) and retrospenial cortex (RSC); 27.4% of total divergences).

In mode 1, we identified 12 HC hits as indicative for family history of ADRD, with the strongest subregion effects identified in CA1, CA2/3, molecular layer, and granule cell layer of the dentate gyrus (DG) (66.7% of HC divergences in mode 1). The remaining HC hits for mode 1 were either located in the parasubiculum, CA4 or hippocampus tail (Fig. 1). We revealed 34 concomitant DN hits, most of them located in the prefrontal cortex (dmPFC, and vlPFC) and posterior midline structures (RSC, PCC, and PCu) which represented 55.9% and 35.3% of total DN hits in mode 1, respectively. As for mode 2, 80.0% of the 10 identified HC hits were located in the left hemisphere (Extended Data Fig. 1). Of those hits, the strongest weights were found in the presubiculum and CA2/3. The remaining HC hits were identified in the CA1, CA4, hippocampal fissure, and DG. While the majority of the 30 DN divergences for mode 2 were located in the prefrontal cortices (dmPFC; 30.0%) and posterior midline structures (PCC and RSC; 26.6%), a substantial proportion of hits were located in the temporal and posterior cortices. In particular, 23.3% of DN divergences for mode 2 were located in the temporal cortices (superior temporal sulcus (STS), middle temporal sulcus (MTS), and temporal pole) compared to 20.0% to the left posterior cortex (inferior parietal lobule (IPL) and superior parietal lobule (SPL)). Mode 3 in turn showed 3 statistically relevant HC hits to the fornix’s fimbria and presubiculum, in concordance with 56 DN divergences (Fig. 2). Of the DN hits identified for mode 3, 35.7% were located in the frontal lobe (dmPFC, vmPFC, vlPFC, pre-supplementary motor area (Pre-SMA), and orbitofrontal cortex (OFC)), 30.3% to posterior midline structures (PCC, RSC, and PCu), 17.9% to the temporal cortices (STS, MTS, and superior temporal gyrus (STG)), and 16.1% to the parietal cortices (IPL, SPL, and temporo-parietal junction (TPJ)). A minority of the modes only showed HC hits, which were either located in the fimbria (mode 8; Fig. 3) or to the hippocampus-amygdala transition area (modes 6 and 10; Extended Data Fig. 2 & 3) without any concomitant DN hits. Inversely, some modes only showed DN divergences in the absence of HC hits. This was the case for mode 4 for which we identified 4 DN hits in the dmPFC (Extended Data Fig. 4), mode 7 for which 9 DN hits were identified in the PFC (dmPFC, and OFC; Extended Data Fig. 5), mode 11 for which 1 DN hit was identified in the PCC (Extended Data Fig. 6), and mode 13 for which 1 DN hit was identified in the STS (Extended Data Fig. 7).

**Figure 1.**
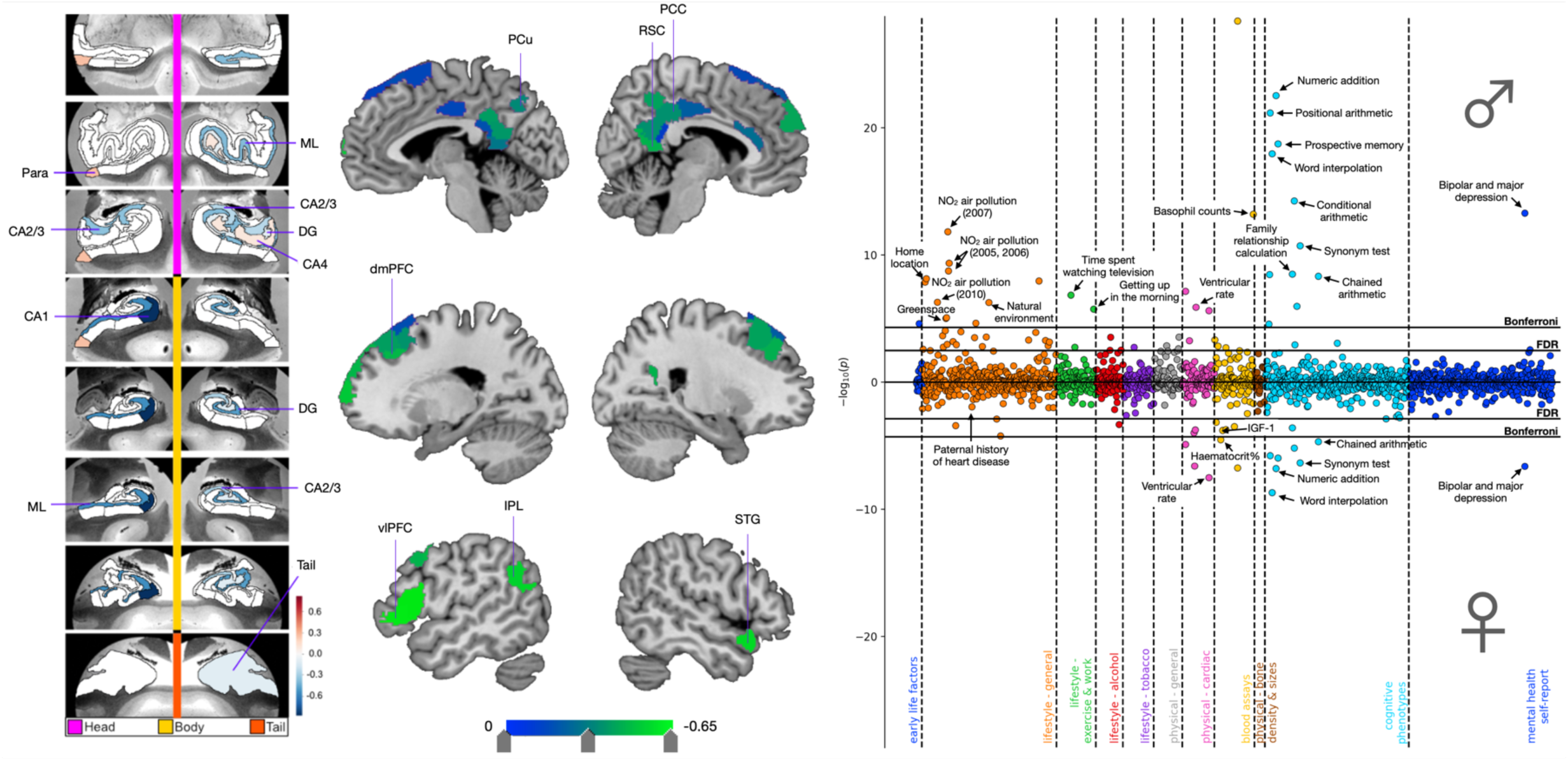
Cognitive, environmental, and cardiovascular phenotypes show sex-specific associations with *APOE* dosage in the context of mode 1. Principled signatures of structural co-variation between 38 hippocampus (HC) and 91 default network (DN) subregions were computed using canonical correlation analysis. 25 hierarchically ordered signatures of HC-DN co-variation, termed *modes*, were derived from this procedure with the most explanatory signature captured by mode 1. A bivariate *APOE* dosage scale was created by summing up positive ‘ɛ2’ and negative ‘ɛ4’ alleles, such that ɛ2 homozygotes would have a dosage of +2 and ɛ4 homozygotes one of -2. The neutral ɛ3 allele was scored as 0. In separate analyses for males (N=17,561) and females (N=19,730), *APOE* dosage was regressed on HC and DN co-variation patterns from mode 1. We then used these sex-specific models to predict *APOE* dosage based on inter-individual expressions of mode 2. *APOE* dosage predicted for each individual was then correlated to 977 UKB phenotypes in separate analyses for males and females. The leftmost and central panels display structural divergences in the HC and DN, respectively, on mode 1 for the group difference analysis of ADRD family history. We identified 12 HC hits mostly located in the cornu amonis (CA) subfields and molecular layer. We also showed 34 DN hits, most of them located in the prefrontal cortex and midline structures. The rightmost panel displays the Miami plot for the correlations between predicted *APOE* dosage in the context of mode 1 and UKB traits. The upper and lower part of the Miami plot displays the correlations for males and females, respectively. The y axis indicates negative decimal logarithms for the p-values of each correlation represented by a dot. We highlight important brain-behaviour associations between *APOE* dosage pooled across subject-specific expressions of mode 1 and verbal-numerical reasoning, supplemented by male-specific correlations with environmental phenotypes. Females showed a specific profile of brain-behaviour associations with cardiovascular phenotypes (e.g., systolic & diastolic blood pressure, insulin-like growth factor 1 (IGF-1), and urea) that extended beyond physical traits shared with males (e.g., cardio-respiratory fitness, and ventricular & pulse rate). ML = molecular layer, Para = parasubiculum, DG = granule cell layer of the dentate gyrus, PCu = precuneus, RSC = retrosplenial cortex, PCC = posterior cingulate cortex, dmPFC = dorsomedial prefrontal cortex, vlPFC = ventromedial prefrontal cortex, IPL = inferior parietal lobule, STG = superior temporal gyrus, FDR = false discovery rate correction.

**Figure 2.**
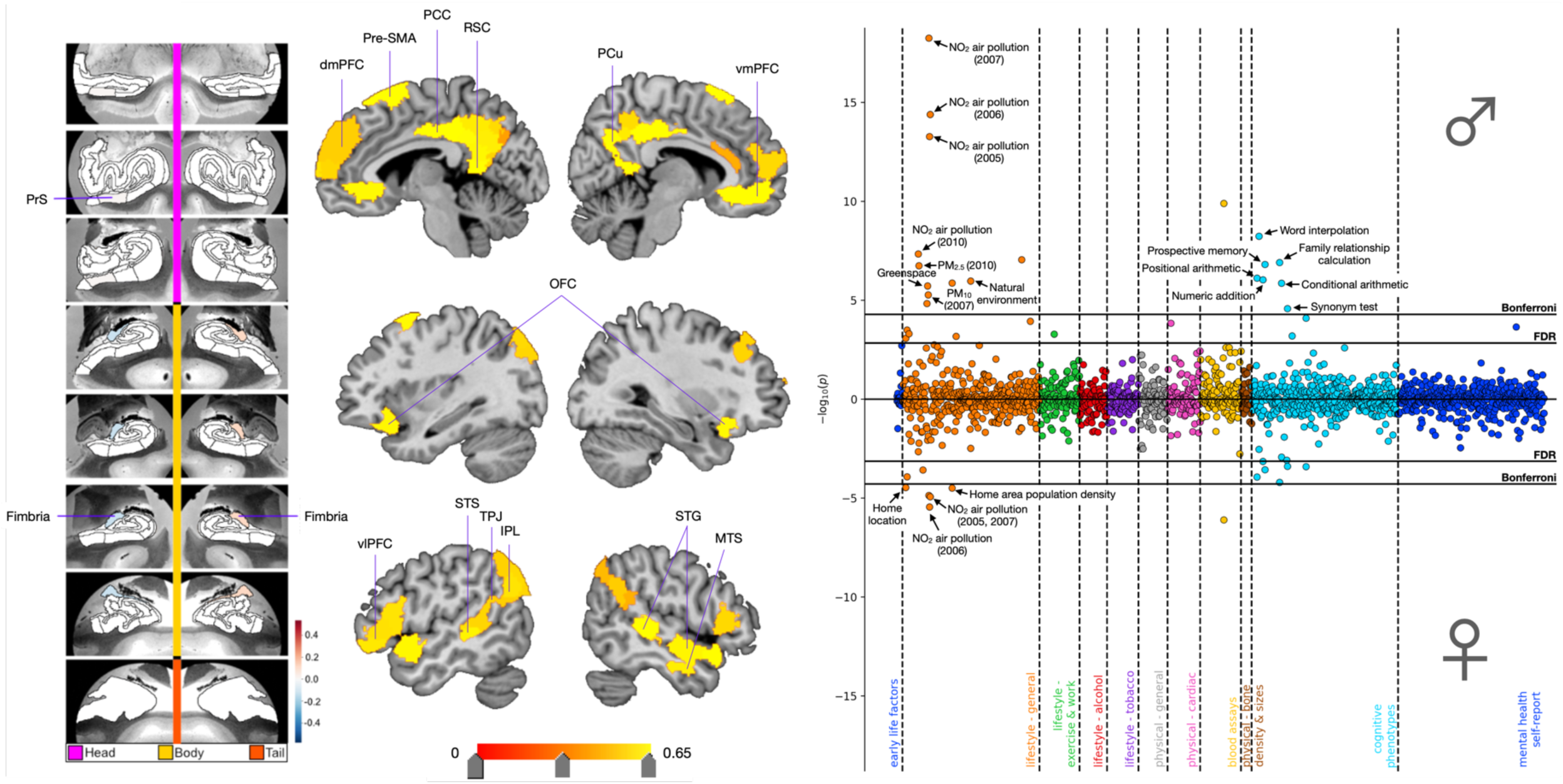
*APOE*-modulated associations for mode 3 revealed a prominence of cognitive and environmental phenotypes in males. Shown here are ADRD-related subregion divergences for mode 3 for the HC (leftmost panel) and DN (central panel). We identified focalized hits to the fimbria and presubiculum with corresponding grey matter differences across the whole DN. In males and females separately, we regressed *APOE* dosage on HC and DN co- variation patterns from mode 3. We then used these sex-specific models to predict *APOE* dosage based on inter-individual expressions of mode 3. The rightmost panel displays the Miami plot for the correlations between *APOE* scores in the context of mode 3 and the portfolio of UKB phenotypes for males (upper half) and females (lower half). We highlighted significant associations with environmental phenotypes, that were again more prominent in males than females. We additionally showed significant correlations with sub-questions of the fluid intelligence battery that were male-specific. PrS = presubiculum, dmPFC = dorsomedial prefrontal cortex, Pre-SMA = pre-supplementary motor area, PCC = posterior cingulate cortex, RSC = retrosplenial cortex, PCu = precuneus, vmPFC = ventromedial prefrontal cortex, OFC = orbitofrontal cortex, vlPFC = ventrolateral prefrontal cortex, STS = superior temporal sulcus, TPJ = temporo-parietal junction, IPL = inferior parietal lobe, STG = superior temporal gyrus, MTS = middle temporal sulcus, FDR = false discovery rate correction.

**Figure 3.**
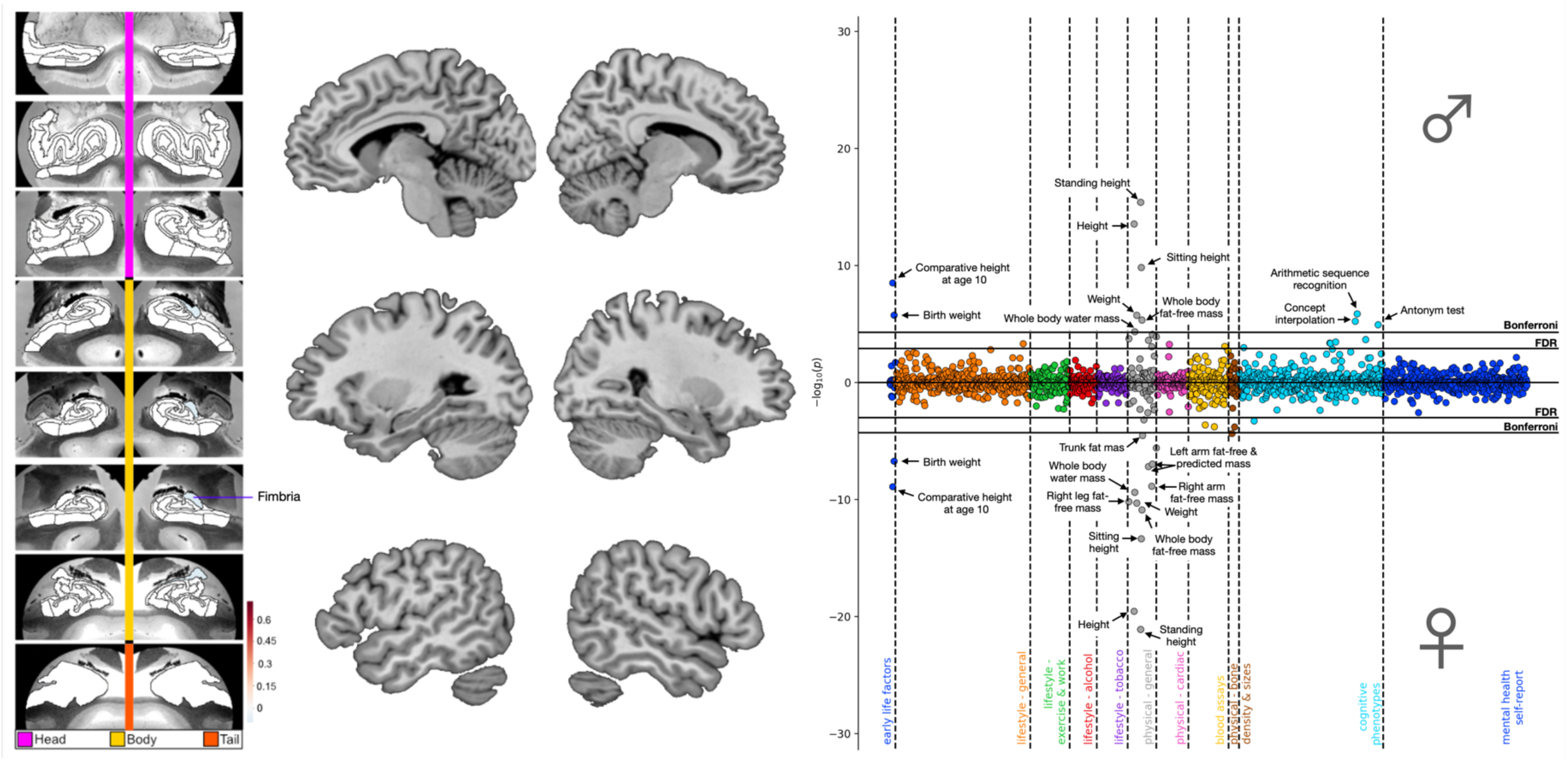
*APOE*-modulated associations for mode 8 linked lipid metabolism to deviation of the fimbria. Shown here are ADRD-related subregion divergences for mode 8 for the HC (leftmost panel) and DN (central panel). We identified a focalized divergence to the fimbria with no corresponding DN hits. In males and females separately, we regressed *APOE* dosage on HC and DN co-variation patterns from mode 8. We then used these sex-specific models to predict *APOE* dosage based on inter-individual expressions of mode 8. The rightmost panel displays the Miami plot for the correlations between *APOE* scores in the context of mode 8 and the portfolio of UKB phenotypes for males (upper half) and females (lower half). We show association with phenotypes related to lipid metabolism and height, supplemented by male-specific associations with sub-questions from the fluid intelligence battery. FDR = false discovery rate correction.

Across HC-DN co-variation signatures, we noted a prominence of HC structural deviation in the CA1, CA2/3 and fimbria for the group analysis of ADRD risk. As for the DN divergences, we highlighted a constellation of structural deviations involving the prefrontal cortices and posterior midline structures. Modes 1 and 2 showed the highest relative numbers of HC divergences (i.e., 12 and 10 hits respectively) as compared to any other modes. While the third signature of HC-DN co-variation only showed 3 statistically relevant HC hits, it showed the highest relative number of DN divergences. Together with mode 8, the focalized divergences found in the fimbria for mode 3 highlighted the importance of the fornix in ADRD risk. We further uncovered concomitant structural divergences in HC and DN subregions with known direct anatomical connections in macaque monkeys such as the presubiculum with RSC [16], and molecular layer with OFC/vmPFC [17]. Ultimately, we revealed an intertwined collection of structural divergences in highly coupled HC and DN subregions which have been linked to ADRD risk and progression by previous research such as the CA1, CA2/3, presubiculum, and the fornix’s fimbria [13, 25–28], as well as the dlPFC, OFC, PCC, and PCu [29–32].

### Phenome-wide fingerprints of brain-behaviour associations uncover sex-specificity in ADRD risk

We next conducted a phenome-wide analysis to systematize domains of UKB traits in terms of their association with HC-DN signatures and ADRD risk. To quantify genetic risk, we created a bivariate dosage scale that tested for the opposing effects of *APOE* ɛ2, often suspected to confer protective benefits [33], and ɛ4, classically believed to escalate dementia risk [4]. We fitted linear regression models to relate inter-individual expressions of HC-DN co-variation from the 25 signatures to subject-level *APOE* ɛ2 vs ɛ4 dosage. Using these linear models, subject-level *APOE* dosage was predicted from a collection of HC-DN signatures and put to the test against 977 curated cUKB phenotypes in a phenome-wide assay, conducted separately in males and females. Only the top three modes with the most brain-behaviour associations across sexes, i.e., mode 1, 3, and 8, are presented below (Figs. 1-3). The phenome-wide profiles for each of the remaining modes with statistically defensible deviations with respect to family history of ADRD are available as part of the online extended data (Extended Data Figs. 1-7).

The phenome-wide profile for mode 1 highlighted brain-behaviour associations with cognitive traits in addition to male-specific correlations with environmental phenotypes (Fig. 1). After carrying out Bonferroni’s correction for multiple comparisons, *APOE* dosage pooled across subject-specific expressions of mode 1 yielded 31 and 13 significant associations in males and females, respectively. Cognitive traits represented 35.5% of significant mode-trait associations in males and 53.8% of those identified in females. Baseline cognitive performance on the fluid intelligence battery accounted for most of the cognitive associations, with 7 questions yielding significant associations in males compared to 6 in females. Significant associations with baseline prospective memory were also identified for both sexes, measured as correct recalling of the object previously shown to participants on the screen. The phenome-wide profiles for both sexes also included ventricular rate on electrocardiogram measured at rest, completion status of electrocardiogram during exercise, and bipolar and major depression status. At the more lenient FDR correction, we observed additional phenotypes linked with erythrocytes count for both sexes. The second most dominant sets of associations for mode 1 centered on environmental phenotypes, such as NO2 exposure, natural environment, and greenspace, which represented 29.0% of significant mode-trait correlations identified in males. Other male-specific associations included lifestyle (time spent watching television and difficulty waking up in the morning) and physiological (hand grip strength, arm mass, and height) phenotypes. At the more lenient FDR correction, males showed additional brain-behaviour associations which included exposure to particulate matter of 2.5 μm and 10 μm or less in diameter (PM2.5 and PM10). After applying Bonferroni’s correction, females showed unique associations with diastolic blood pressure and hematocrit percentage. When applying FDR correction, additional cardiovascular phenotypes showed significant associations in females such as paternal history of heart attack, systolic blood pressure, insulin-like growth factor 1 (IGF-1), and haemoglobin concentration. In sum, our phenotypical profiling assay highlighted important phenome-wide associations between *APOE* dosage pooled across subject-specific expressions of mode 1 and verbal-numerical reasoning, supplemented by male-specific correlations with environmental phenotypes. Females instead showed a specific profile of brain-behaviour associations with cardiovascular phenotypes that extended beyond physical traits shared with males.

In the phenome-wide profile for mode 3, we uncovered brain-behaviour associations with cognitive and environmental phenotypes, again more prominent in males than females (Fig. 2). After Bonferroni’s correction, *APOE* dosage in the context of mode 3 expressions yielded 19 and 6 significant mode-trait associations in males and females, respectively. Environmental phenotypes represented 52.6% of significant associations in males and 83.3% of those identified in females. Significant associations with NO2 exposure and home area population density were observed for both sexes. Males also showed significant associations with baseline cognitive performance on 6 questions from the fluid intelligence battery as well as with baseline prospective memory. Females did not show significant associations beyond those shared with males, with the exception of home location. At the more lenient FDR correction, females showed additional associations with prospective memory and baseline cognitive performance on 5 questions from the fluid intelligence battery. As such, *APOE* dosage pooled across subject-specific expressions of mode 3 allowed us to uncover a rich portfolio of associations with environmental and cognitive phenotypes that were more robust in males than females.

In comparison to the overlapping portfolio of brain-behaviour associations derived from mode 1 and 3, the phenome-wide profile for mode 8 emphasized a unique set of physiological phenotypes (Fig. 3). After Bonferroni’s correction, *APOE* dosage pooled across subject-specific expressions of mode 8 yielded 11 and 15 significant mode-trait associations in males and females, respectively. Physical phenotypes related to body mass and height represented 55.5% of significant correlations in males and 80.0% of those identified in females. After Bonferroni’s correction, males showed significant associations with cognitive performance on 3 questions from the fluid intelligence battery assessed in the online follow-up. At the more lenient FDR correction, males showed further associations with cognitive performance on 2 additional questions from the fluid intelligence battery and with the maximum number of digits remembered correctly on the numeric memory test, both assessed in the online follow-up. After Bonferroni’s correction, females showed significant associations with trunk fat mass and heel bone mineral density. In sum, we highlighted important phenome-wide associations between *APOE* dosage pooled across subject-specific expressions of mode 8 and proxies of cardiovascular health, supplemented by male-specific correlations with cognitive phenotypes. The phenome-wide profiles derived across these three concomitant regimes of HC-DN co-variation emphasized sex differences in ADRD risk, with recurring associations with air pollution and verbal-numerical reasoning that were more prominent in males than females.

### APOE gene variants are associated with distinct clusters of risk-anatomy links

We next examined ADRD-specific clusters of risk-anatomy links across each unique *APOE* gene variant (i.e., ɛ2/2, ɛ2/3, ɛ3/3, ɛ2/4, ɛ3/4, and ɛ4/4). We computed the interactions between the subject-specific expressions of HC-DN covariation modes (canonical variates) and each *APOE* genotype (encoded as binary variables, such that participants who do not carry a given genotype were zeroed out). In doing so, we obtained six new population-wide indices, that is, one for each distinct *APOE* genotype that we correlated, using Spearman’s coefficient, with 63 curated ADRD risk factors (a phenotype collection used previously [34]). We then performed an agglomerative clustering analysis which consisted in a nested merging of correlation coefficients with similar variance until all observations got merged in a single cluster. The ensuing dendrograms indicated the distance between each cluster identified when retaining three levels of branching (Fig. 4).

**Figure 4.**
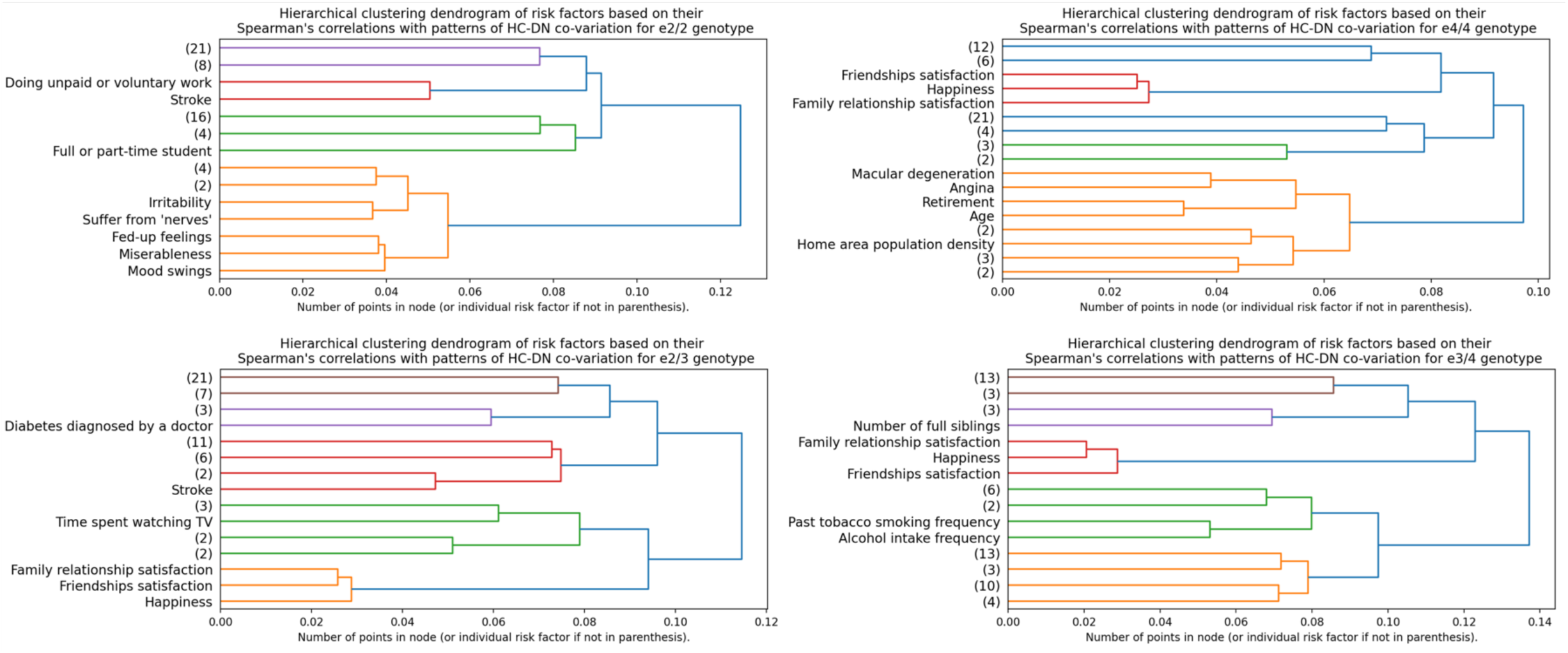
Social, personality, and age-related phenotypes show risk-anatomy links across *APOE* gene variants. To test for *APOE*-modulated brain-behaviour interactions, we multiplied the population-wide HC and DN co-variation patterns by each *APOE* genotype coded as binary variables such that participants who do not carry a given genotype were zeroed out. In doing so, we obtained six new population-wide vectors, i.e., one for each distinct *APOE* gene variant: ɛ2/2, ɛ2/3, ɛ3/4, ɛ4/4, ɛ3/3 (Extended Data Fig. 8), and ɛ2/4 (Extended Data Fig. 8). We then computed the Spearman’s correlations between these six new vectors and the 63 pre-selected Alzheimer’s disease risk factors to test for risk-anatomy links. We performed an agglomerative clustering analysis on these Spearman’s correlations, which consists in repeatedly merging Spearman’s correlations with similar variance together until all observations are merged into a single cluster. Here are shown the dendrograms which indicate the distance between each cluster identified when retaining three levels of branching for *APOE* ɛ2/2 (upper left; N=217), ɛ2/3 (lower left; N=4,625), ɛ4/4 (upper right; N=822), ɛ3/4 (lower right; N=8,613). We showed the early emergence of a social cluster that comprises ‘family relationship satisfaction’, ‘happiness’, and ‘friendships satisfaction’ for most *APOE* gene variants except from ɛ2/2. In the dendrogram for ɛ2 homozygotes, we rather observed the early branching of neuroticism- related phenotypes. As for the dendrogram for ɛ4 homozygotes, we also observed the early branching of age-related phenotypes, such as macular degeneration, angina, retirement, and participant age. While social phenotypes formed important risk-anatomy links in most *APOE* genotypes, we found that neuroticism was uniquely associated with ɛ2 homozygotes, while age-related phenotypes were prominent to ɛ4 homozygotes.

Our integrated analysis of risk-anatomy links showed the relatively early branching of a social cluster for ɛ2/2, ɛ2/3, ɛ3/4, ɛ4/4, ɛ3/3, and ɛ2/4 (Extended Data Fig. 8), genotypes that comprised ‘happiness’, ‘family relationship satisfaction’, and ‘friendships satisfaction’. The presence of this social cluster across most *APOE* gene variants suggests that these three self-reported social phenotypes show similar risk-anatomy links with HC-DN patterns independently from ɛ4 dosage. While this social cluster was apparent in individuals with ɛ2/3 genotype, it was not clearly observable amongst ɛ2 homozygotes at the same level of branching. Rather, we noted the early emergence of a personality cluster that comprised self-reported traits related to neuroticism, such as miserableness, irritability, and mood swings. The specificity of this personality cluster suggests that neuroticism is an important phenotype that is interlocked with HC-DN co-variation patterns in ɛ2 homozygotes. In the clustering analysis for ɛ2 homozygotes, we also showed the emergence of self-reported phenotypes related to social engagement, such as ‘doing unpaid or voluntary work’ and being a ‘full or part-time student’. The neuroticism and social engagement clusters were located on opposing branches of the dendrogram for ɛ2 homozygotes which possibly reflects their diverging relations with the disclosed HC-DN signatures. In ɛ4/4 individuals, in addition to the social cluster, we observed the early branching of age-related phenotypes, such as macular degeneration, angina, and retirement. Aside from emphasizing the importance of social factors in individual expressions of HC-DN co-variation, we uncovered different constellations of risk-anatomy links specific to ɛ2 and ɛ4 homozygotes. We revealed that neuroticism, which is known to be closely linked to loneliness [35], is a personality trait specific to ɛ2 homozygotes as reflected by unique patterns of association with HC-DN co- variation expressions. In contrast, the clustering analysis for ɛ4 homozygotes was characterized by age-related diseases that have previously been linked to lower socioeconomic status [36–38]. In sum, we have characterized clusters of risk-anatomy links that were either shared by most *APOE* gene variants (e.g., social phenotypes) or unique to ɛ2 and ɛ4 homozygotes.

### Sex-specific dependencies between APOE gene variants and signatures of HC-DN co-variation explain ADRD risk

We next directed attention to sex-specific interactions between HC-DN covariation regimes and *APOE* genotype status that would explain inter-individual differences in ADRD risk. To this end, we tested whether HC-DN signatures systematically interacted with specific *APOE* genotypes in explaining variation in a collection of 63 ADRD risk factors (cf. above). More formally, each risk factor was individually regressed on the subject-specific expressions of HC and DN patterns for each of the 25 modes. This analysis step hence supplied 50 estimated linear models per target risk factor. Each model treated as input variables the main effect of the HC or DN pattern expressions, the main effects of the six *APOE* genotypes, and the interaction between each *APOE* genotype with the HC or DN pattern, controlling for age. Separate analyses were carried out in the male (Fig. 5, leftmost panels) and female (Fig. 5, rightmost panels) subgroups of our UKB cohort. To ascertain the robustness of our findings, we compared each coefficient estimate against empirically data-derived null distributions obtained through a rigorous permutation procedure (i.e., label shuffling permutation). We only interpreted the model coefficients that emerged as statistically relevant based on the respective null distributions at 95% confidence.

**Figure 5.**
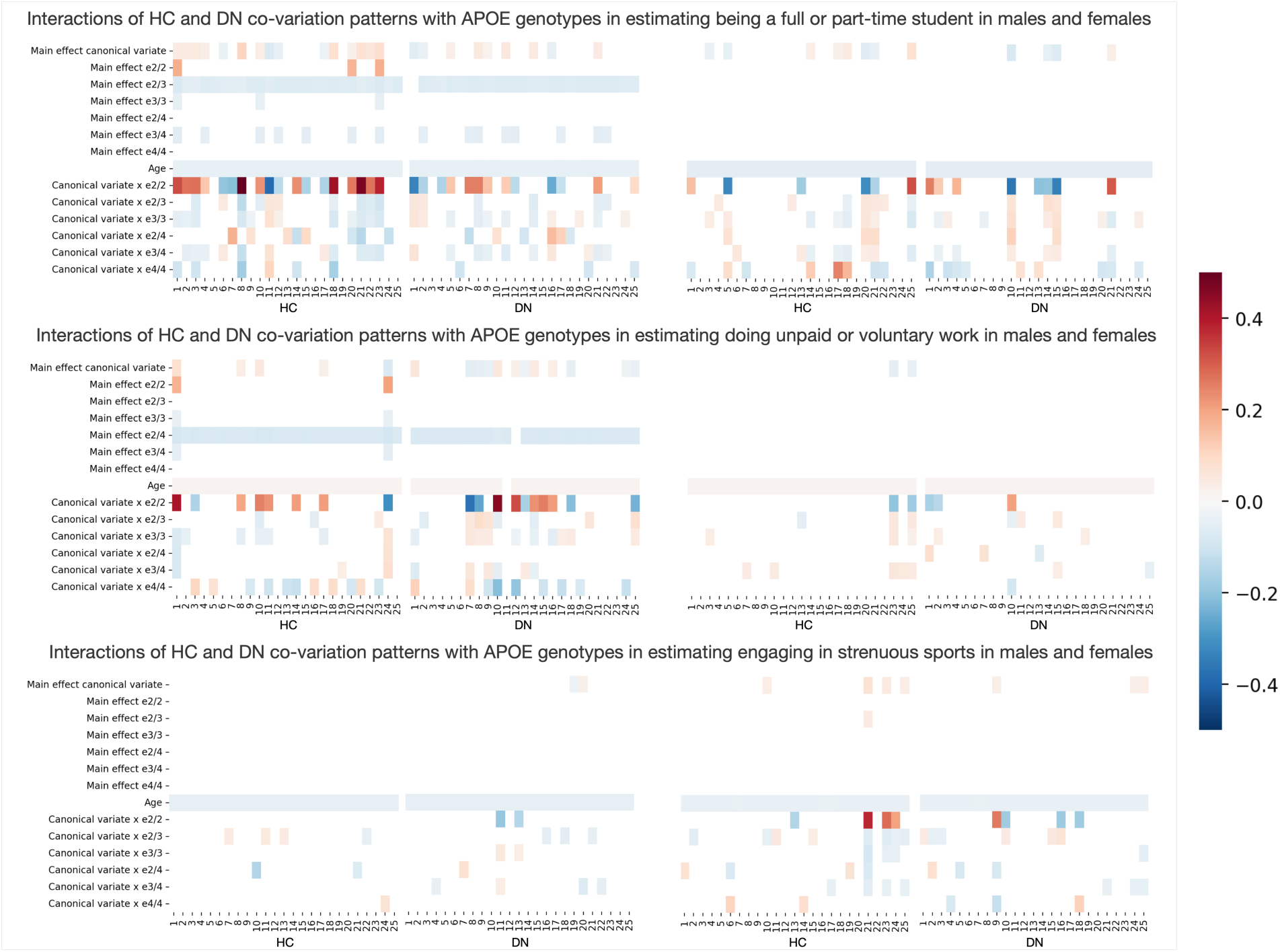
Brain-*APOE* ɛ2/2 interaction explains variance in social lifestyle in males and physical activity in females. We tested whether HC-DN signatures interacted with *APOE* genotypes in explaining variance on the 63 pre-selected ADRD risk factors. Each risk factor was individually regressed on a single HC or DN pattern from a given mode of HC- DN co-variation, resulting in 50 different linear models per risk factor. Each model took as parameters the main effect of a given HC or DN pattern, the main effects of the six *APOE* genotypes (i.e., ɛ2/2, ɛ2/3, ɛ3/3, ɛ2/4, ɛ3/4, and ɛ4/4), and the interaction between each of the six *APOE* genotype and the given HC or DN pattern, controlling for age. Separate analyses were run for males (leftmost plots) and females (rightmost plots). Each column on the heat maps thus represents the coefficients for a single linear regression model. The first 25 columns show the coefficients for HC patterns, whereas the last 25 columns show the coefficients for DN patterns. We assessed the robustness of our findings by comparing each coefficient to empirically built null distributions obtained through permutation testing. Only the coefficients that were statistically different from their respective null distributions in 95% of the time are presented. We displayed the modifiable risk factors for which the strongest brain-APOE interactions were observed. In the top panels, we show that *APOE* ɛ2/2 interact with HC and DN co-variations patterns in estimating being a full or part-time student, with stronger coefficients observed for males on the HC side (regression models 1- 25). Similarly, on the middle panels, we show that *APOE* ɛ2/2 preferentially interact with HC and DN canonical variates in estimating doing unpaid or voluntary work in males. In the bottom panels, we show that *APOE* ɛ2/2 interact with selective HC and DN canonical variates in estimating engaging in strenuous sport in females. We have thus showed that *APOE* ɛ2/2 interacts preferentially with HC-DN co-variation patterns in estimating social lifestyle in males, and physical activity in females. These interactions profiles suggest that ɛ2, and not ɛ4, is driving most of the brain-genes interactions in healthy individuals at risk of developing ADRD with a substantial level of sex- specificity.

Across a comprehensive set of analyses across 63 ADRD risk factors, we identified the strongest interaction effects in homozygote ɛ2 carriers. Notably, brain-*APOE* interactions accounted for more variance in several modifiable social and cardiovascular risk factors than did the main effects of *APOE* ɛ2 and ɛ4. Across both sexes, ɛ2 homozygotes showed strong interaction with HC and DN patterns for full vs part-time student. Male ɛ2 homozygotes showed strong interactions with HC and DN pattern expressions for doing unpaid or voluntary work. In parallel, female ɛ2 homozygotes showed strong interactions with HC-DN patterns expressions for engagement in strenuous sports. The only exception to this trend was observed for our examination of the physical environment score – a measure of environmental factors, such as air quality, air emission, and proximity to waste disposal or industrial sites. This risk factor led to strong brain-*APOE* interactions in male and female homozygote ɛ4 carriers (Extended Data Fig. 9). Across the different domains of risk factors investigated, we singled out brain-*APOE* interactions specific to ɛ2 and ɛ4 homozygotes that were not identifiable in heterozygotes and non-carriers. While we observed no appreciable sex effect for the interaction of *APOE* ɛ4/4 and HC-DN co-variation expressions, we found defensible sex-specificity for the role of *APOE* ɛ2/2. More precisely, we showed strong interactions between *APOE* ɛ2/2 and HC-DN co-variation patterns for social lifestyle factors in males and physical activity factors in females. Through our analyses of a variety of risk factors, we have thus isolated brain-*APOE* interactions unique to ɛ2 and ɛ4 carriers that differed in their level of sex-specificity.

After examining target risk factors, we next put to the test whether expressions of HC-DN signatures bear relations with *APOE* genotypes in explaining ADRD risk. In dedicated analyses for males (Fig. 6, upper panels) and females (Fig. 6, lower panels), family history of ADRD was regressed on a single HC or DN pattern, resulting in 50 different linear models per sex. Each such model was fed as input variables the main effect of the HC or DN pattern, the main effects of the *APOE* genotypes, and the interactions between each *APOE* genotype and the HC or DN pattern, controlling for age. We assessed the robustness of our findings by comparing each coefficient to empirically built null distributions obtained through permutation testing (cf. above). We focused interpretation on the model coefficients that were statistically robust against their respective null distributions at 95% confidence. We found no statistically relevant main effect of *APOE* ɛ2/2 on ADRD risk amongst males. For *APOE* ɛ2/3 and ɛ3/3 carriers, in turn, we found similar effects on ADRD risk in males, lowering the odds of ADRD family history by approximately 30% across the different models investigated. Likewise, *APOE* ɛ2/4 and ɛ3/4 carriers showed similar effects in tracking ADRD risk in males, elevating the odds of ADRD family history by more than 20% on average. As expected from the literature, *APOE* ɛ4/4 increased the odds of ADRD family history by more than 56% in males across the different models investigated. In females, *APOE* ɛ2/2 status decreased the odds of ADRD family history by 50% on average, while ɛ2/3 and ɛ3/3 status led to decreases of approximately 25% and 17%, respectively. In contrast, *APOE* ɛ3/4 and ɛ4/4 status lifted the odds of ADRD family history by approximately 35% and 86%, respectively. Among females, *APOE* ɛ2/4 carriers were associated with dampened ADRD risk relative to *APOE* ɛ3/4 carriers. The odds of ADRD family history associated with *APOE* ɛ2/4 were only increased by 24% in females. This ∼10% reduction in ADRD risk, uniquely observed amongst females, could be taken to suggest that ɛ2 can still be protective against ADRD risk in the presence of an ɛ4 allele. Females also showed some strong brain-*APOE* interactions above and beyond the well-established risk and protective effects associated with each *APOE* genotype. Notably, the interaction of mode 9 DN pattern expressions with *APOE* ɛ2/2 status was associated with a 2-fold increase in ADRD risk. It was considerably stronger than the main risk effect conferred by *APOE* ɛ4/4. This strong interaction effect can be taken to suggest that HC-DN co-variation plays a chief role in ADRD risk, which might have been overlooked by previous analyses restricted to genetic data. In sum, we identified and annotated sex-specificity in the opposing effects of ɛ2 and ɛ4 on ADRD risk, with demonstrably stronger brain-*APOE* interactions amongst females.

**Figure 6.**
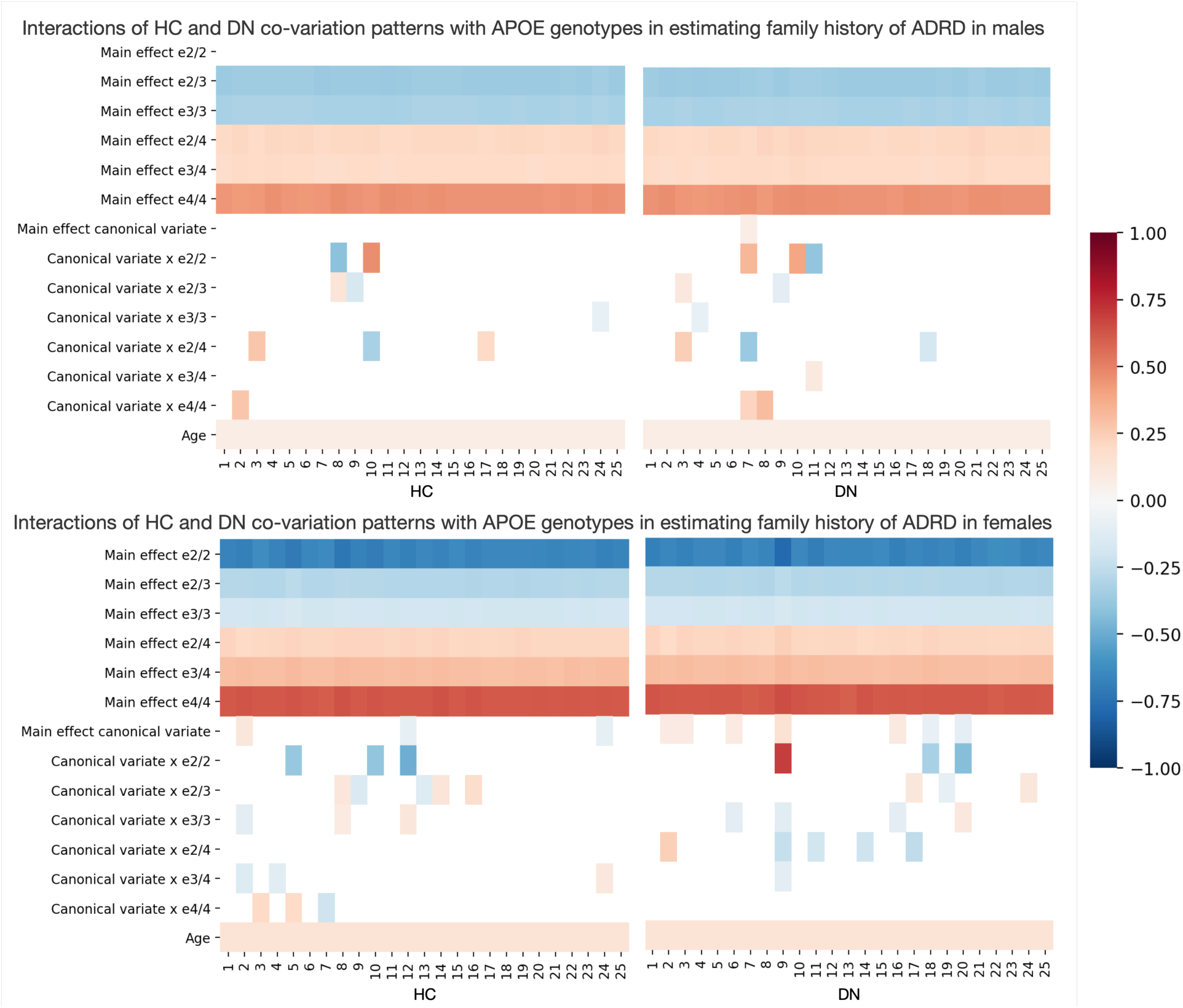
The protectiveness of ɛ2 is sex-dependant and modulated by HC-DN co-variation patterns. In separate analyses for males and females, we tested whether HC-DN signatures interacted with *APOE* genotypes in explaining variance on ADRD risk. Family history of ADRD was regressed on a single HC or DN pattern, resulting in 50 different linear models per sex that each took as parameters the main effect of the HC or DN pattern, the main effects of the *APOE* genotypes, and the interactions between each *APOE* genotype and the HC or DN pattern, controlling for age. Separate analyses were run for males (higher plots) and females (lower plots). Each column on the heat maps thus represents the coefficients for a single linear regression model. The first 25 columns show the coefficients for HC patterns, whereas the last 25 columns show the coefficients for DN patterns. We assessed the robustness of our findings by comparing each coefficient to empirically built null distributions obtained through permutation testing. Only the coefficients that were statistically different from their respective null distributions in 95% of the time are presented. We found that the main effect of *APOE* ɛ2/2 against ADRD risk was only statistically robust in females. We also showed a spectrum in the opposing effects of ɛ2 and ɛ4 amongst females, such that ɛ2/4 was associated with a lower increase in ADRD risk than did *APOE* ɛ3/4, which in turn was associated with a lesser risk than ɛ4/4. We further found that the protectiveness of *APOE* ɛ2/2 interacts with brain structure and can even lead to increase in ADRD risk amongst females with a strong expression of mode 9. These interactions profiles suggest that the protectiveness of ɛ2/2 is not only sex-specific, but also modulated by HC-DN co-variation expressions.

### Dominant principles of brain-behaviour associations uncovered a male-specific link with neuroticism

In a final suite of analyses, we conducted an exploratory principal component analysis (PCA) to disentangle the major sources of brain-behaviour variation in our UKB cohort. We first computed Pearson’s correlations between the 25 pairs of expressions (i.e., canonical variates) from the HC and those from the DN patterns and the 63 pre-selected ADRD risk factors. This step yielded 3,150 distinct coefficients represented by a risk by canonical variates matrix (X63 x 50). We then carried out a PCA to reduce the dimensionality to three major axes of brain-behaviour. These explained ∼16.3%, ∼12.8%, and ∼8.2% of the total variance in the cross-correlation matrix, respectively (Extended Data Fig. 10).

The leading axis of variation highlighted the previously identified social cluster (happiness, family relationship satisfaction, and friendships satisfaction) along other socioeconomic (number of vehicles in household, number of people in household, and physical environment score) and cardiovascular risk factors (stroke, hypertension, diabetes). The second most important axis mainly emphasized socioeconomic and lifestyle factors (fluid intelligence, education, alcohol intake frequency, average household income, and time spent watching TV). The third most explanatory axis tracked neuroticism and its associated personality trait indicators (miserableness, fed-up feelings, mood swings, worrier/anxious, sensitivity/hurt feelings, and worry too long after embarrassment) from the rest of the risk factors. We again emphasized the importance of social factors on HC-DN co-variation expressions along other cardiovascular, cognitive, and personality phenotypes highlighted by previous analyses.

To certify the robustness of our findings, we performed a split-half reliability assessment of our principal component solution. We derived two random participant subsets of equal size (N=18,645) from the original UKB sample; and repeated the analogous procedure described above. The axes obtained from the first random subset had explained variances of ∼13.9%, 11.1%, and 7.9%, respectively, whereas those obtained from the second subset had explained variances of ∼15.3%, 11.2%, and 9.1%, respectively. The three major axes of brain-behaviour associations derived from each subset of participants yielded very similar summary statistics and flagged the same sets of risk factors emphasized along each corresponding dimension (Supp. Fig. 1). Hence, our reliability assessment indicated that the PCA solution was robust.

We then repeated the identical pattern-learning workflow sex-stratified in males and females separately. The three major axes of brain-behaviour associations derived from each group were roughly the same as in our full-sample PCA solution; except for the third axis (Supp. Fig. 2). While neuroticism was highlighted by the latter brain-behaviour association axis amongst males, no such association was observed for females. The third axes of brain-behaviour association for females rather emphasized social (looking after one’s home or family, attending religious group) and cardiovascular (e.g., engagement in strenuous sport, sleep duration) risk factors. As expected from these results, the first two axes explained similar amount of variance in the data in males and females (∼14.6% and ∼11.9%) while sex-specific divergence in explained variance was observed for the third axis (∼9.6% in males compared to ∼7.4% in females). We thus concluded that the association between neuroticism-related phenotypes and HC-DN co-variation expressions was mainly male-specific. As such, our sex-specific analysis revealed that the two major axes of brain-associations were shared across sexes, whereas the third axis emphasized sexually dimorphic groups of risk factors.

## Discussion

Longstanding research has insisted on alteration of the DN and HC in early ADRD development (see for example [14]). However, brain-imaging investigations seldom had the opportunity to incorporate rare genotypes such as *APOE* ɛ2/2. At the same time, common epidemiological studies that have reported the protective effect of carrying an ɛ2 allele are not typically equipped to perform an adequately powered brain-imaging examination at the scale of thousands of people. We overcame several shortcomings by capitalizing on *APOE* genotyping and structural brain scans from ∼40,000 UK Biobank participants. Our mission-tailored analytical framework was especially designed for disentangling ADRD-specific differences in brain structure at the population level. Revisiting ADRD through this lens, we uncovered sex-specific associations between rarely investigated *APOE* gene variants and microstructurally defined HC-DN signatures hardly ever discerned in a prospective human cohort. Our collective findings paint a more concrete picture of the antagonistic effects of *APOE* ɛ2 and ɛ4 on population-wide HC-DN signatures, along with their interlocking divergences between men and women.

Epidemiological studies, without access to brain-imaging assessments, have provided evidence suggesting that an ɛ2 allele typically acts to protect against late onset Alzheimer’s disease [22, 33] and against Aβ accumulation [39–43]. The protective qualities of ɛ2 status have been noted even in the presence of an ɛ4 allele [12]. Nonetheless, sex-specific impact of *APOE,* especially its ɛ2 gene variants, on brain structure could seldom be investigated at the population level. By deriving an envelope of distinct HC-DN signatures at a fine-grained resolution amongst thousands of healthy adults, we were able to uncover brain-*APOE* interactions systematically overlooked by traditional brain-imaging studies. Stratifying our population cohort by sex and *APOE* gene variants, we showed that the protective effect of *APOE* ɛ2/2 on ADRD risk was not statistically robust amongst males, even in a sample of ∼20,000 participants. In contrast, we demonstrated a spectrum of ɛ2 and ɛ4 effects amongst females such that *APOE* ɛ2/4 was associated with milder ADRD risk than ɛ3/4, which in turn was associated with milder ADRD risk than ɛ4/4. Resilience towards cognitive decline generally observed amongst ɛ2 carriers could arise from relatively higher baseline APOE steady state levels in regions including the HC and frontal cortex as compared to ɛ4 carriers and ɛ3 homozygotes [44–47]. Isoform specific effects related to the APOE protein could be further enhanced by microglia-driven homeostatic responses to Aβ accumulation [48, 49]. Older ɛ2 carriers with amyloid pathology, who are biologically more efficient at scavenging the peptide, are indeed less likely to be diagnosed with dementia than ɛ3 homozygotes of the same age [50]. Cell proliferation and survival in the HC is thought to be particularly modulated by estrogens [51–53] which could have downstream impact on microglial and astrocytic APOE synthesis [54]. The presence of an estrogen-dependent enhancer in the promoter region of the *APOE* gene is thus bound to favour female ɛ2 carriers [55]. Theses previous elements of evidence are in line with our present finding that the protective effect of *APOE* ɛ2 on ADRD risk is sex-specific and also unique to specific HC-DN co-variation patterns. Notably, we found that female ɛ2 homozygotes with a high expression of mode 9 had twice the odds of having family history of ADRD. We have thus shed light on important nuances in the predominant genetic account of ADRD by questioning the protectiveness of ɛ2 when placed in relation to sex and brain structure.

We extended the discovered sex differences in ADRD risk by highlighting a female-specific constellation of brain-behaviour associations with cardiovascular traits. As the neuroprotective effect of estrogen weakens in older ages, women become more vulnerable to neurovascular disorders that can ultimately lead to dementia [56]. Cardiovascular risk factors that are exacerbated in females following menopause, such as trunk fat mass, have been associated with chronic neuroinflammation and microstructural alteration of the fornix [57, 58] – the main output tract from the HC that carries direct neural signals towards partner regions of the midline DN [59]. Building on existing literature, we identified ADRD-related divergences in the fimbria of the fornix in healthy participants for mode 8 that showed selective brain-behaviour associations with proxies of cardiovascular health (e.g., water mass, fat-free mass, and weight). For the same HC-DN signature, we found a female-specific association with trunk fat mass, a correlate of estrogen declines [60]. This observation supports a link between cardiovascular health, female sex, and microstructural alteration of the fornix. Despite the protective effect of *APOE* ɛ2 against ADRD previously discussed, carrying an ɛ2 allele has been associated with elevated risks for cardio- and neurovascular disorders [61–65]. *APOE* ɛ2 is indeed limited in its ability to mediate the vascular clearance of cholesterol metabolites and triglycerides which could in turn precipitate the risks of cholesterol pathologies such as hyperlipoproteinemia and cardiovascular sequelae [66]. The variability of the protective effect of physical activity on dementia risk when stratifying participants by ɛ4 status might be taken to suggest that *APOE* ɛ2 is driving the relationship between physical activity and cognitive performance [67–71]. Engaging in physical activity could therefore be particularly beneficial to older female ɛ2 carriers in counteracting the rising risk of neurovascular complications resulting from the combined effect of *APOE* ɛ2 and decreased estrogen levels. Bringing support for this claim, we have shown specific interactions between HC-DN signatures and *APOE* ɛ2/2 genotype in explaining variation in physical activity – an effect that we found exclusive to females. The specificity of this effect to ɛ2 homozygotes is consistent with previous findings that have associated ɛ2 to increased longevity in centenarians [72]. Given that almost 90% of centenarians are females, the sex-specificity of our results is consistent with a genotype-driven behavior that favors longevity via exercise in female ɛ2 homozygotes.

Cardiovascular risk factors are known to interact with *APOE* in older age. For example, converging evidence now suggests that ɛ2 carriers are at greater risks of macular degeneration [63] and atherosclerosis [66]. Nevertheless, we observed the early branching of cardiovascular phenotypes such as angina and macular degeneration on the clustering analysis for ɛ4 homozygotes. While this seems to run counter previous evidence, the presence of ‘retirement’ and ‘home area population density’ within the same cluster as cardiovascular phenotypes may suggest an interplay between socioeconomic status and ADRD co-morbidities. Since ɛ4 carriers are known to be especially vulnerable to the detrimental effects of air pollution on cognition [73–80], residence in urban areas, which is characteristic of lower socioeconomic status, could exacerbate the risks of developing dementia and other co-morbidities. Elevated risks of angina, ischemia, and coronary heart disease have indeed been observed amongst individuals with lower socioeconomic status [36, 37], even after retirement [38]. In a sample of UKB participants, socioeconomic status was previously associated with grey matter volume in subregions of the DN [81]. Here, we have shown ADRD-related divergences such as the dmPFC and TPJ. The brain-behaviour associations we observed for HC-DN signatures and cardiovascular phenotypes could indirectly arise from environmental contributions that depend on socioeconomic status. This non-linear effect could in turn be accentuated by an ɛ4-specific vulnerability to pollution-induced neurotoxicity. As such, we have found unique associations between HC-DN signatures and *APOE* ɛ4/4 in explaining variance in physical environment score, a composite measure of air quality, air emission, and proximity to waste disposal or industrial sites.

Indeed, epidemiological studies have provided evidence that traffic-related air pollution and residence near major roadways are associated with decreased cognitive abilities [77, 82–89] and higher risk of developing dementia [78, 90–98]. Our phenome-wide assay tied mode 1 expressions to blood markers (e.g., erythrocytes, hemoglobin, and haematocrit) and air pollution. This supports an interplay between environmental stressors, vascular integrity, and dementia. Mode 1 also showed 19 DN hits in the PFC – a subregion in which vascular and perivascular white matter damage has been specifically observed in humans and canines chronically exposed to high levels of air pollutants [99]. Such accumulation of nanoscale particulate matter in endothelium cells, basement membranes, axons, and dendrites coincided with prefrontal white matter damage; which is in line with deficits in the blood-brain barrier [99]. Autopsy samples from patients with Alzheimer’s disease have further shown reduced pericyte coverage in CA1 and PFC (Brodmann area 9/10). These were two subregions in which we showed ADRD-related structural divergences in mode 1, as compared to healthy blood vessels in controls [100]. We have thus identified subregions that are consistent with early vascular leakage in the aging brain, such as CA1 and PFC, as manifesting ADRD-related structural deviation in the same HC-DN signature associated with air pollution in our phenome-wide analysis. In doing so, we extend on the plausible role of vascular integrity in protecting the brain from environmental stressors that might precipitate ADRD onset.

In a similar vein, in-vitro analyses have suggested that exposure to air pollution can trigger microglial activation which in turn can cause oxidative stress [101, 102]. Pollution-triggered oxidative stress could be particularly detrimental to males as they are thought to display lower expression of antioxidant enzymes responsible for scavenging reactive oxygen species [103, 104]. As a result, male mice show up to 4-fold higher rates of oxidative toxicity in astrocytes, neurons, and mitochondria compared to female mice [103, 105]. Building on these experimental findings primarily from rodent species, we found that the association between HC-DN co-variation and air pollution was male-specific. Parts of the DN are thought to be amongst the earliest sites of Aβ accumulation [29] and to consume some of the highest oxygen levels in the entire brain [106]. As such, the DN sticks out as a hotspot for both oxidative stress and ADRD pathology. A previous study has indeed found widespread glucose hypometabolism in the DN of ADRD patients that was associated with increased levels of CSF lactate, a marker of mitochondrial damage, in the OFC and mPFC as compared to cognitively healthy controls [107]. Recent evidence suggesting that Aβ1-42 acts on reactive oxygen species to induce glucose hypometabolism [108]. Hence, the combined effect of air pollution and amyloid pathology could be particularly detrimental in exacerbating ADRD risk amongst males. In line with an effect on escalating ADRD risk, specifically in males, we have linked ADRD-related structural deviation in the OFC and mPFC with a profile of associations with environmental phenotypes for mode 3. As was the case for mode 1, these associations were more prominent in males than females. In addition to emphasising a male-specific vulnerability to neurotoxicity, our phenome-wide analysis pointed towards a female-specific resilience to pollution-mediated impairment and subsequent neuronal death. For example, our phenome-wide profile for mode 1 derived, for females, did not show statistically relevant associations with air pollutants but displayed a significant correlation with IGF-1. Estrogen and IGF-1 are thought to exert synergetic, non-additive effects on neurite outgrowth and survival; presumably by acting on a single neuroendocrine pathway [109]. IGF-1 is secreted by neurons and glia and possibly acts as a neurotrophic factor regulating neuroendocrine function in the central nervous system [109]. Subcutaneous injection of IGF-1 has previously been associated with increased neurogenesis in the adult rat brain [110, 111]. In mode 1, in addition to a female-specific association with IGF-1, we have shown HC hits in the granule cell layer of the DG and in CA4, which are two subfields in which neurogenesis has been observed in rodents [110, 112, 113] and primates [114]. Together with its associated divergences in HC-DN co-variation expressions, the phenome-wide profile for mode 1 shed light on a female-specific resilience towards pollution-induced impairment and subsequent neuronal death. While scarcely reported in human subjects, these sex-specific divergences in vulnerability to neurotoxicity — observed here for both mode 1 and 3 — are hence in accordance with experimental findings from animal models.

Building on the knowledge that ADRD and verbal-numerical reasoning share overlap in underlying genetic architecture [115], we showed significant brain-behaviour associations between ADRD risk and baseline cognitive performance on the fluid intelligence battery for top modes 1, 2, and 3. While previous investigations of fluid intelligence and ADRD in the UKB were often limited to genetic evidence [115–118], we highlighted distinct HC-DN signatures related to verbal-numerical reasoning at the population level. In doing so, we found prominent ADRD-related structural divergences in the left CA1, CA2/3, presubiculum, and fimbria, which are amongst the first and notorious regions to be affected by ADRD pathology [13, 25–27]. Some authors have claimed that white matter disruption may trigger grey matter degradation in the HC and higher-order neocortex by activating a maladaptive neuroinflammatory response [119]. Changes in fornix microstructure have indeed been reported in individuals at risks of ADRD before the onset of clinical symptoms [26] and subsequently identified as an accurate predictor of progression from mild cognitive impairment to ADRD [27]. Consistent with an early involvement of the fornix in ADRD-associated cognitive deficits, we showed structural divergence in the fornix’s fimbria and 56 DN regions for mode 3, which were accompanied by a profile of associations with questions from the fluid intelligence battery.

Recent brain-imaging evidence has extended the concept of a hippocampally mediated cognitive map to interpersonal relationships by highlighting the involvement of the DN, and hence the fornix, in schematic representations of the self and others. Notably, fMRI results from Tavares and colleagues suggest that the HC tracks how we represent others in a social hierarchy while the PCC/PCu, key hubs of the DN, tracks the social distance between ourselves and others [120]. Consistent with a reliance on the HC-DN pathway for human-defining aspects of spatiotemporal processing, we found a brain-behaviour association with navigating family relationship, a subtest of the fluid intelligence battery, that was significant in males for mode 3.

We have thus provided a link between verbal-numerical reasoning and ADRD risk that was accompanied by alterations in HC and DN subregion covariation regimes involved in episodic processing.

By exploring risk-anatomy links across the different *APOE* gene variants, we have tied the subjective appraisal of one’s social bonds to subject-specific expressions of HC-DN co-variation signatures. In addition to reinforcing the joint implication of HC and DN subregions in navigating social relationships, we built on previous UKB work that has linked population regimes of HC-DN co-variation to loneliness, a subjective indicator of social deprivation [24]. Zajner and colleagues have indeed linked loneliness to midline structures of the DN (e.g., RSC and dmPFC) and HC (e.g., CA1 and molecular layer) in which we have shown ADRD-related hits for mode 1 [24]. In concordance with these previous results that have highlighted the importance of loneliness to HC-DN co-variation expressions, our clustering analysis of risk-anatomy linked especially uncovered a distinct set of social phenotypes. This social cluster, that related to happiness, family relationship satisfaction, and friendships satisfaction, emerged across most *APOE* gene variants in ways that were independent of ɛ4 status and departed from traditional ADRD risk factors. Loneliness has previously been associated with a faster rate of cognitive decline, especially within subcomponents of episodic memory sustained by the DN such as delayed and immediate recall [121–124]. In older age, a decrease in social activity possibly related to unemployment and/or retirement could increase feelings of loneliness and consequently escalate the risk of cognitive decline and ADRD [125]. Social disengagement has indeed been associated with the incidence of cognitive decline amongst older adults [126–128]. In contrast, engaging in social activities has been linked with up to a 40% decrease in ADRD risk [67, 126, 129]. While social support has been associated with a dampened stress response [130], loneliness is thought to affect not only neuroendocrine but also immune functions [131, 132]. The faster rate of cognitive decline observed amongst lonely individuals is thus in line with evidence suggesting that neuroinflammation exacerbates ADRD pathology [133]. Volunteering and having student status, two social engagements that have repeatedly been flagged in our analyses on ɛ2 homozygotes, could possibly counteract the pathological stress response in lonely older adults. Our study has thus uncovered risk-anatomy links that are consistent with the involvement of social factors as preventing or exacerbating ADRD risk. We have especially linked the subjective appraisal of one’s social ties to HC-DN co-variation expressions amongst most *APOE* genotypes, along some associations with measures of social engagement that were specific to ɛ2 homozygotes.

Our collective findings further indicate neuroticism to be an important phenotype with unique ties to HC-DN co-variation expressions in ɛ2 homozygotes. Neuroticism, which is intimately related with loneliness [35], could predispose individuals to ADRD by by weakening strong social support ties and increasing chronic stress through dysregulation of the hypothalamus-pituitary-adrenal (HPA) axis [131, 134]. In fact, the HC subiculum, presubiculum, and parasubiculum are believed to have direct connections to the hypothalamus via the fornix [135]. These connections could provide a pathway through which the subjective appraisal of one’s relationships, which can in turn results in loneliness or neuroticism if social needs are unfulfilled, is conveyed to the HPA to affect the stress response. Prospective cohort studies have indeed linked neuroticism to higher risks of developing cognitive impairments [136] and dementia [137–139]. Yet, no effects of ɛ4 dosage on cognitive decline has been observed in neurotic individuals in these previous reports [136, 138]. The absence of relationship between *APOE* and neurotic traits reported by previous studies might arise from restricting analyses to ɛ4 carriers [136, 138]. Indeed, the combined analysis of ɛ4 and the K variant of BCHE, another genetic risk factor associated with ADRD, revealed an intriguing association between the combined risk alleles, increased basal levels of serum glucocorticoids, cognitive performance and lower self-esteem in older adults [140]. The ramifications of neuroticism for ADRD risk, which might be underscored by *APOE* ɛ2, have been overlooked in all these studies. A prolonged state of neuroticism-related metabolic imbalance could be especially detrimental to ɛ2 homozygotes in precipitating the development of neurovascular disorders typically observed in older age [56, 66]. Consistent with this hypothesis, recent evidence has shown that having a positive outlook on aging, such as a sense of purpose, amplified the protective effect of *APOE* ɛ2 against cognitive decline [141]. The protective effect of *APOE* ɛ2 on cognition was enhanced for individuals with positive beliefs on aging and reduced for those with negative beliefs to the point where ɛ2 carriers no longer showed a significant cognitive advantage [141]. Our results add elements to this literature by suggesting that having a negative outcome on life, which is characteristic of a neurotic personality type, is especially detrimental to ɛ2 carriers as reflected by unique patterns of brain-behaviours associations with specific HC and DN subregions. The opposing health effects of neuroticism and social activity are possibly reflected by the brain, as social and neurotic phenotypes were divided in two groups when clustered based on their correlation with HC-DN covariation regimes for ɛ2 homozygotes. Our study thus reinforces the detrimental effect of neuroticism on ADRD risk and characterized its unique interplay with HC-DN co-variation expressions in ɛ2 homozygotes.

In sum, the typically protective benefits conferred by *APOE* ɛ2 regarding ADRD risk have mainly been discussed in epidemiological cohort that were not designed to incorporate inter-individual differences in high-resolution brain structure assessments. In contrast, neuroimaging investigations of healthy participants before the onset of ADRD-associated clinical symptoms have focused on characterizing the functional correlates of ɛ4 carriership. Our present study has reconciled these two approaches by contrasting profiles of brain-behaviours associations characteristic of *APOE* ɛ2 and ɛ4 in a large epidemiological cohort of ∼40,000 participants. In doing so, we were uniquely position to illuminate sex-specific associations with modifiable risk factors that were unique to ɛ2 and ɛ4 homozygotes. Key risk factors relevant to ɛ2 carriers included neuroticism, social disengagement, and physical inactivity. In contrast, environmental phenotypes that repeatedly emerged in our results as being linked to ADRD risk were characteristic of ɛ4 homozygotes. These distinct risk factors could guide potential clinical interventions and governmental policies.

## Author contribution statement

All authors designed the study, analyzed and interpreted the results, and wrote the manuscript.

## Online methods

### Population data source

The UK Biobank (UKB) is a large-scale data-collection initiative that offers in-depth information on ∼500,000 participants recruited from across Great Britain (https://www.ukbiobank.ac.uk/). This rich epidemiological cohort comprises a wide variety of resources including physical and cognitive assessments, as well as demographic and health records. In addition to the availability of genetic data for most participants through a genotyping array (and more recently through whole-exome sequencing), the UKB provides multi-modal imaging scans that are routinely augmented and will extend to ∼100,000 participants by the end of 2022. The present study was based on the data release from February/March 2020. To ensure reproducibility, we adopted the uniform preprocessing pipelines designed and carried out by FMRIB, Oxford University, UK [142]. Building on a uniform quality-control workflow enables a better comparison to other and future UKB research. At the time of data release, expert-curated image-derived phenotypes of grey matter morphology (T1-weighted magnetic resonance imaging) were available for 38,292 participants. Grey matter phenotypes from these participants were used to compute dominant regimes of structural correspondence between the HC and DN and identify anatomical subregions that systematically differentiate individuals with and without family history of ADRD. As for all subsequent analysis steps, we focused on the 37,291 participants with both *APOE* single nucleotide polymorphisms (SNP) genotyping (rs429358 and rs7412) and brain-imaging measures (47% men and 53% women). These participants were aged 40-70 years when recruited (mean age 54.8, standard deviation [SD] 7.5 years). The present analyses were conducted under UK Biobank application number 25163. Further information on the consent procedure can be found elsewhere (http://biobank.ctsu.ox.ac.uk/crystal/field.cgi?id=200).

### Target phenotype for ADRD risk

We used self-reported family history of ADRD as a simple but accurately measurable estimate of ADRD risk. Maternal (UKB data field 20110) and paternal (UKB data field 20107) history of ADRD was ascertained as part of the initial assessment (2006-2010). There was a total of 9,776 (25.5%) participants with self-reported parental history of ADRD within the brain-imaging cohort of 38,292 participants. Of those with family risk, 6,820 UKB participants reported an occurrence of ADRD on their mother’s side and 3,675 participants on their father’s side. A minority of participants reported both maternal and paternal history of ADRD (719 individuals).

Most genome-wide association studies have adopted a case-control framework that focused on the difference in allele frequency between patients with ADRD and healthy controls [143, 144]. While useful in identifying risk loci associated with clinical diagnosis, this approach might not be best suited to derive a reliable estimate of ADRD liability in the general population. When dealing with late-onset diseases, such as ADRD, using ‘proxy cases’, that is, the relatives of affected individuals, could allow for a more complete characterization of disease risk amongst individuals before the onset of clinical symptoms [145]. It was a key advantage that working with proxy cases also allowed to boost the sample size and thus the statistical power of our quantitative analyses to identify more suitable effects. In particular, self-reports of family history of ADRD in the UKB, precisely the same phenotype at the core of the present investigation, was found to replicate established risk loci from case-control investigations as well as identify novel loci [145, 146].

### Brain-imaging and preprocessing procedures

Magnetic resonance imaging (MRI) scanners (3T Siemens Skyra) were matched at several dedicated data collection sites with the same acquisition protocols and standard Siemens 32- channel radiofrequency receiver head coils. Brain-imaging data were defaced, and any sensitive meta-information was removed to protect the anonymity of the study participants. Automated processing and quality control pipelines were deployed [142, 147]. To improve homogeneity of the imaging data, noise was removed by means of 190 sensitivity features. This approach allowed for the reliable identification and exclusion of problematic brain scans, such as due to excessive head motion.

The structural MRI data were acquired as high-resolution T1-weighted images of brain anatomy using a 3D MPRAGE sequence at 1 mm isotropic resolution. Preprocessing included gradient distortion correction (GDC), field of view reduction using the Brain Extraction Tool [148] and FLIRT [149, 150], as well as non-linear registration to MNI152 standard space at 1 mm resolution using FNIRT [151]. To avoid unnecessary interpolation, all image transformations were estimated, combined, and applied by a single interpolation step. Tissue-type segmentation into cerebrospinal fluid, grey matter, and white matter was applied using FAST (FMRIB’s Automated Segmentation Tool, [152]) to generate full bias-field-corrected images. In turn, SIENAX [153] was used to derive volumetric measures normalized for head sizes.

Sub-segmentation of the DN was anatomically guided by the Schaefer-Yeo reference atlas [154]. We extracted a total of 400 parcels among the 7 canonical networks, 91 of which defined as belonging to the DN. Volume extraction for 38 HC subregions was conducted using Freesurfer automatic sub-segmentation [21] which drew on an ultra-high resolution (∼0.1mm isotropic) probabilistic atlas. As part of the Freesurfer 7.0 suite, HC sub-segmentation was refined by carefully considering surrounding anatomical structures.

As a preliminary procedure, these MRI-derived measures were cleaned to remove inter-individual variation in brain region volumes that could be explained by nuisance variables. Building on previous UK Biobank research [155, 156], we regressed out the following variables of no interest from each brain-derived volume measure: body mass index, head size, head motion during task-related brain scans, head motion during task-unrelated brain scans, head position and receiver coil in the scanner (x, y, and z), position of scanner table, as well as the data acquisition site, in addition to age, age^2^, sex, sex*age, and sex*age^2^. Sex was acquired from the National Health Service (NHS) central registry and updated by the participant if incorrect (UKB data field 31). The nuisance-cleaned volumetric measures served as the basis of our primary co- decomposition analysis – seeking to quantify how the 91 DN subregions co-deviate with the 38 HC subregions in the context of ADRD risk.

### Population co-variation between hippocampus subregions and default-network subregions

At the heart of our analysis workflow, we derived dominant regimes of structural correspondence that provide insights into *how structural variation among the finely segregated HC can track structural variation among the finely segregated DN*. We employed canonical correlation analysis (CCA), a doubly multivariate statistical technique, to identify population “signatures” of HC-DN co-variation. CCA was a natural choice of method as it is especially designed to disentangle patterns of joint correlation between two high-dimensional variable sets

[157–159]. The first variable set, *X*, was constructed from subject-level grey matter volume in DN subregions (number of participants x 91 DN parcels matrix). The second variable set, *Y*, was constructed from HC subregion volumes (number of participants x 38 HC parcels matrix). The two variable sets can be formally described as followed:

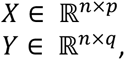

where *n* denotes the number of observations or UKB participants, *p* is the number of DN subregions, and *q* is the number of HC subregions. Subregion volumes from both variable sets were z-scored across participants to zero mean (i.e., centering) and unit variance (i.e., rescaling).

CCA then addressed the problem of maximizing the linear correlation between low-rank projections from two variable sets or data matrices [23]. The two sets of linear combinations of the original variables are obtained by optimizing the following target function:

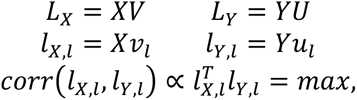

where *V* and *U* denote the respective contributions of *X* and *Y*, *L*_*X*_ and *L*_*Y*_ denote the respective latent ‘modes’ expression of joint variation (i.e., canonical variates) based on patterns derived from *X* and patterns derived from *Y*, *l*_*X,l*_ is the *l*th column of *L*_%_, and *l*_*Y,l*_ is the *l*th column of *L*_&_.

Our CCA application thus sought to identify linear combinations of *X* and *Y* that optimize their low-rank projections in the derived latent embedding. Such an approach resulted in pairs of latent vectors with subject-specific expressions *l*_*X,l*_ and *l*_*Y,l*_ (i.e., canonical variates) with maximized joint correlation. Corresponding pairs of latent vectors were found by iteratively decomposing the data matrices *X* and *Y* into *k* components, where *k* denotes the number of modes given the model specification. In other words, CCA searched for the canonical vectors *u* and *v* that maximizes the symmetric relationship between the data matrices of DN subregion volumes (*X*) and HC subregion volumes (*Y*). In doing so, CCA identified the two concomitant projections *Xv_l_* and *Yu_l_* that optimized the correspondence between structural variation in the segregated DN and HC.

Put differently, each principled signature of HC-DN co-variation, or mode, represents the cross-correlation between a constellation of within-DN volumetric variation and a constellation of within-HC volumetric variation that co-occurred in conjunction with each other. The set of *k* modes are mutually uncorrelated by construction (orthogonality) [158]. They are also naturally rank-ordered based on the amount of variance explained between the embedded allocortical and neocortical volume sets [158]. The first and strongest mode therefore explained the largest fraction of joint variation between (linear) combinations of HC subregions and (linear) combinations of DN subregions. Each ensuing cross-correlation signature captured a fraction of structural variation that is not explained by one of the *k* − 1 other modes. The Pearson’s correlation between a pair of canonical variates (i.e., canonical correlation) is commonly used to quantify the linear correspondence between HC subregions and DN subregions for a given mode. The two variable sets were entered into CCA after a confound-removal procedure based on previous UK Biobank research (cf. above).

### Group difference analysis

After constructing population signatures of conjoint HC-DN co-variation, we performed a rigorous group difference analysis to single out microstructural divergences in specific anatomical subregions with respect to ADRD family history. For each of the derived modes of HC-DN co- variation, we aimed to isolate anatomical subregions that show statistically defensible deviation in individuals with and without family history of ADRD. To do so, we carried out a principled test that assessed any statistically relevant differences in the solution vector obtained from the CCA (i.e., canonical vectors, cf. above) of individuals at ADRD risk compared to the control group without ADRD family history (cf. above for target phenotype).

Following previous UK Biobank research [24, 160], we robustly characterized the difference between individuals with and without family history of ADRD by carrying out a bootstrap difference test of the CCA solution at hand [161]. The goal of this approach was to identify consistent patterns of deviation that differentiate subjects with and without family history of ADRD. We first proceeded by constructing several alternative datasets that we could have gotten (with the same sample size) which capture the underlying population variation. For each of the 100 bootstrap iterations, these alternative datasets were built by randomly pulling participant samples with replacement. In each such bootstrap iteration, we estimated two CCA models in parallel by fitting one separate model to each of the two groups. In doing so, we carried out 2 * 100 separate model estimations of the doubly multivariate correspondence between HC subregions and DN subregions.

To compare the CCA solution in individuals with and without family history of ADRD, we matched corresponding modes based on sign invariance and mode rank order. Canonical vectors of a given mode that carried opposite signs were aligned by multiplying one with -1. The importance rank of the CCA modes was adjusted by sorting Pearson’s correlation coefficients between pairs of corresponding canonical vectors (i.e., canonical correlations) from strongest to weakest. To estimate a quantity of group difference in relation to ADRD risk, we performed the elementwise subtraction of the corresponding canonical vector entries of a given mode *k* between the two groups. Pooling outcomes across the 100 bootstrap iterations, we thus aggregated the difference estimate for each canonical vector entry, thereby quantifying the uncertainty deviation for each particular HC or DN subregion.

By probing the underlying population variation, we were able to quantify the degree of uncertainty within each of our derived modes of HC-DN co-variation. For each identified population signature, we therefore isolated statistically defensible group differences in microanatomically defined HC and DN subregions. ADRD-related structural divergences were determined by whether the two-sided confidence interval included zero or not according to the 10/90% bootstrap-derived distribution of difference estimates [156]. In doing so, we obtained a non-parametric estimate of how ADRD risk is manifested in specific subregions for each of the 25 examined HC-DN signatures.

### SNP genotyping: six variants of APOE gene

We capitalized on our large sample size to demystify the HC-DN co-variation expressions associated with ɛ2 allele and ɛ4 allele homozygotes, compared to their heterozygous counterparts for the ɛ2, ɛ3 or ɛ4 alleles. Genotype-level sampling and quality control procedures for the UKB are available online (https://biobank.ndph.ox.ac.uk/showcase/label.cgi?id=263). *APOE* genotypes were determined based on two SNPs: rs429358 and rs7412. *APOE* ɛ4 was determined as the combination of rs429358(C) and rs7412(C). *APOE* ɛ2 was determined as the combination of rs429358(T) and rs7412(T). *APOE* ɛ3 was determined based on rs429358(T) and rs7412(C). A total of 37,291 participants had both *APOE* genotyping and brain-imaging-derived measures. Among those participants, 9,525 (25.5%) reported family history of ADRD. We observed 6 different *APOE* gene variants in our population sample: ɛ3/3 (59.3%), ɛ3/4 (23.1%), ɛ2/3 (12.4%), ɛ2/4 (2.4%), ɛ4/4 (2.2%), and ɛ2/2 (0.6%), which correspond to frequencies expected from a population primarily composed of people from European decent [22]. Contrasting the effect of ɛ2 vs ɛ4 allele dosage on inter-individual expressions of HC-DN co- variation enabled us to quantify the degree to which distinct *APOE* allelic combinations are characteristic of ADRD risk (cf. next section). In doing so, we aimed to interrogate gradual dosage effects in brain-*APOE* associations, rather than simply look at ɛ4 carrier vs non-carrier status.

### Phenome-wide analysis of brain-behaviour associations in relation to ɛ2 vs ɛ4 dosage

We performed a rich annotation of the HC-DN co-variation signatures by means of their phenome-wide association with UKB traits. We were interested in how ɛ2 vs ɛ4 allele dosage is manifested in inter-individual expressions of HC-DN co-variation, and how these manifestations in turn relate to UKB traits amongst a variety of predefined risk categories. We benefited from a rich portfolio of phenotypes encompassing lifestyle, cognitive, mental, and physical health assessments to ascribe profiles of brain-behaviour associations to each of the 25 modes of HC-DN co-variation.

We started with a raw collection of ∼15,000 phenotypes that we fed into the FMRIB UKB Normalisation, Parsing And Cleaning Kit (FUNPACK version 2.5.0; https://zenodo.org/record/4762700#.YQrpui2caJ8). FUNPACK was used to extract phenotype information covering 11 major categories, including cognitive and physiological assessments, physical and mental health records, blood assays, as well as sociodemographic and lifestyle factors. We removed any brain-imaging-derived information. The diet category was additionally excluded from downstream analyses as it contained only 4 phenotypes. FUNPACK was designed to perform automatic refinement on the UKB data, which included removing ‘do not know’ responses and fill the blank left by unanswered sub-questions. For example, the amount of alcohol drunk on a typical drinking day for a participant who indicated not drinking would be scored as zero drinks, even though this sub-question was not actually asked at assessment. FUNPACK’s output consisted in a collection of 3,330 curated phenotypes which were then fed into PHEnome Scan ANalysis Tool (PHESANT [162], https://github.com/MRCIEU/PHESANT) for further refinement. In addition to data cleaning and normalization, PHESANT categorized the data as belonging to one of four datatypes: categorical ordered, categorical unordered, binary, and numerical. Categorical unordered variables were one-hot encoded, such that each possible response was represented by a binary column (true or false). The final curated inventory comprised 977 phenotypes spanning 11 FUNPACK-defined categories.

We next checked for statistically robust associations between HC-DN signatures and the portfolio of 977 extracted phenotypes with respect to ADRD genetic risk. We used a one-step stacking strategy [163, 164] to predict genetic risk as a function of individual expressions of HC-DN co-variation. Data stacking consists in using a “base” model, often linear regression [164], to express an input vector in a lower-dimensional space. The output of the base model, which often consists in a single variable, can then be used as a single predictor in a new “stacking” model. Data stacking therefore addressed the problem of selecting a single best predictor out of a combination of highly correlating input variables — which in our case were the HC and DN co- variation patterns. Such an approach allowed us to re-express a whole signature of HC-DN co- variation in terms of the degree it tracked the associated risk conferred by *APOE*. We formed a single continuous number that represented how much a given HC-DN signature is reflective of ɛ2 vs ɛ4 dosage for a given individual. For that aim, we created a bivariate dosage scale by summing up positive ‘ɛ2’ and negative ‘ɛ4’ alleles, such that a homozygous individual carrying *APOE* ɛ2/2 would have a score of +2 and one carrying *APOE* ɛ4/4 a score of -2. The neutral *APOE* ɛ3 allele, usually considered as baseline risk in epidemiological studies [22], was scored as 0. Using a bivariate dosage scale made it possible to investigate the antagonistic effects of ɛ2 and ɛ4 in a single model. In doing so, we stayed faithful to our overarching goal of unravelling their adversarial impact on HC-DN co-variation.

Aiming to capture possible sex-specific effects, we regressed the ɛ2 vs ɛ4 dosage on inter- individual expressions of a given mode in males and females separately. We thus estimated 2 * 25 different base models, one for each HC-DN signature and each sex, that each had two parameters: the pair of co-variation expressions (i.e., canonical vectors, cf. above) associated with the HC and DN patterns. We used these 25 regression models to explain the subject-level ɛ2 vs ɛ4 dosage as a function of HC-DN co-variation expressions. For each subject and mode combination, we asked *what would the expected ɛ2 vs ɛ4 dosage be given this subject’s specific expression of HC-DN co-variation?* For each subject, we hence used the regression model to explain a range from -2 to +2 for each mode, which represented the ɛ2 vs ɛ4 dosage associated with their individual expression of HC-DN co-variation. For each mode, we selected the 5^th^ and top 95^th^ percentiles to identify the top 5% and lower 5% individuals who were more vs less likely to develop ADRD based on the derived ɛ2 vs ɛ4 dosage risk. We focused on the extreme of the dosage distribution to target the brain-*APOE* associations especially linked to ɛ2 and ɛ4. The analogous approach is widely adopted in genome-wide analyses to remove associations not directly linked to the target genotype [165, 166].

For each sex separately and for a given mode, the designated participants were put to a test of association with the 977 curated UKB phenotypes, with appropriate correction for multiple comparisons. The Pearson’s correlation between a phenotype and genetic risk predicted based on a specific HC-DN signature revealed both the association strength and accompanying statistical significance of the given mode-trait association. For each HC-DN signature, two widely used procedures were carried to adjust for the multitude of associations being assessed. First, we adjusted for the number of tested phenotypes by using Bonferroni’s correction for multiple comparisons (0.05/977 = 5.11e-5). Second, we used the false discovery rate (FDR), another popular adjustment, although less stringent than Bonferroni’s correction. The false discovery rate [167] was set as 5% [168–170] and computed for each HC-DN signature in accordance with standard practice [171]. For the sake of visualization, we used Miami plots to compare the profiles of brain-behaviour associations derived from males and females. For visualization purposes, phenotypes in Miami plots were coloured and grouped according to the category membership defined by FUNPACK.

#### Clustering of risk factors based on their correlation with HC-DN co-variation expressions

We next systematically explored non-linear associations between established ADRD risk phenotypes and HC-DN co-variation expressions across the different *APOE* gene variants. Our goal was to probe for clusters of risk factors that are interrelated with the derived patterns of HC and DN co-variation. To this end, we used a hierarchical clustering approach that allowed us to assess the relative importance of ensuing clusters in each of the different *APOE* genotypes to explore gradual *APOE* dosage effects on risk-anatomy links.

We adopted a targeted approach by focusing on a set of 63 risk factors (collection of phenotypes used previously [34]), including classical cardiovascular and demographic traits, as well as social richness indicators recently linked to ADRD in the UKB cohort. The first step of the clustering analysis consisted of multiplying the z-scored canonical variates by each of the six one-hot encoded *APOE* genotypes (i.e., ɛ2/2, ɛ2/3, ɛ3/3, ɛ2/4, ɛ3/4, and ɛ4/4) such that participants without a given genotype were zeroed out. The six ensuing matrices (number of participants x 50 canonical variates) represented the individual expressions of HC-DN co-variation signatures for participants with a given *APOE* genotype whereas other participants were scored as 0s. We then computed the Spearman’s correlation between these six genotype-specific matrices and the z-scored risk factor matrix (37,291 participants x 63 risk factors) to investigate risk-anatomy links. Spearman’s correlation is a nonparametric metric of statistical dependence between the rankings of two variables that can be used to capture monotonic non-linear phenomena. The Spearman’s correlation coefficients reduce to the Pearson’s correlation between the rank values of two variables, and hence range from -1 (inversely proportional association) to +1 (proportional association). We obtained a new cross-association matrix *X* ∈ R^*+ - ./^ which represented the Spearman’s correlation between the 63 risk factors and the 50 canonical variates for each of the six *APOE* genotypes. The obtained Spearman’s correlation coefficients thus carried the non-linear association strength of a given risk-anatomy link for a particular *APOE* genotype.

For each of the six *APOE* genotypes, we performed an agglomerative hierarchical clustering analysis on *X* to regroup risk factors based on their 50 associations with HC-DN co- variation pattern expressions. We used Ward’s minimum variance method [172] to compute the linkage matrix between the Spearman’s correlation coefficients of each risk-anatomy link in Euclidian space. Ward’s minimum variance criterion consists in minimizing the total within-cluster variance defined as the error sum of squares:

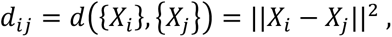

where *d*_*ij*_ represents the squared Euclidean distance between two points (or cluster of points) *i* and *j*. At each step, the pair of coefficients or preceding candidate clusters that gives the minimum increase in within-cluster variance is selected for merging. The procedure was performed recursively until all coefficients were merged into a single cluster. For each of the six *APOE* genotypes, we could thus create a dendrogram that represented the distance in Euclidian space between the clusters retained after three levels of branching. The level of branching refers to the number of divisions from the final merge. The dendrograms allowed us to visualize the clustering results for each of the six *APOE* genotypes at the same level of branching and identify meaningful clusters of risk-anatomy links that are shared or unique.

### Regression of ADRD risk on HC-DN signatures and APOE gene variants

We next tested whether specific *APOE* genotypes showed interaction effects with signatures of HC-DN co-variation in explaining inter-individual differences in ADRD risk. As our goal was to highlight previously overlooked sex effects, we conducted our interaction analyses in males and females separately. In doing so, we aimed to characterize brain-*APOE* interactions in relation to their sex-specific impact on ADRD risk.

A first series of analyses consisted in regressing each of the 63 previously investigated ADRD risk factors on *APOE* genotypes, co-variation patterns from the HC and DN sides (i.e., canonical variates), and the interaction between *APOE* genotypes and co-variation patterns, controlling for age. Aiming to capture possible sex-specific effects, we conducted separate analyses in males and females. Each of the 25 modes of HC-DN co-variation was represented by two regression models: one for its HC pattern and one for its DN pattern. We thus formed 50 univariate regressions models, in males and females, for each of the 63 risk factors. In each of these models, a given risk factor was regressed on one HC or DN canonical variate, the six *APOE* genotypes (ɛ2/2, ɛ2/3, ɛ3/3, ɛ2/4, ɛ3/4, and ɛ4/4), and six interaction terms capturing the non-linear association between each of the six *APOE* genotypes and the given HC or DN pattern, controlling for age. Each regression model thus aimed at explaining variance in one of the 63 risk factors for a given sex based on these 14 parameters.

For each regression model, we performed a rigorous permutation analysis to assess the robustness of each of the 14 regression coefficients. In 1,000 iterations, we randomly shuffled the outcome variable (i.e., a given risk) across participants and recomputed the otherwise identical regression model based on the data with randomized outcome. We recorded the regressions coefficients from each of the 1,000 iterations and used them to build empirical null distributions on which we performed two-tail statistical tests. We considered statistically relevant coefficients that differ from their respective null distributions in at least 95% of the iterations, which ensured that we were at least 5% certain that the effect was robustly different from zero.

A second series of analyses consisted in regressing family history of ADRD on a set of explanatory input variables including i) *APOE* genotypes, ii) co-variation patterns from the HC and DN sides (i.e., canonical variates), and iii) the interaction between *APOE* genotypes and co- variation patterns, controlling for age. For each sex, we built separate logistic models for each of the 25 HC and 25 DN canonical variates, for a total of 50 models per sex. In each model, family history of ADRD (encoded as 0 for no and 1 for yes) was regressed on one HC or DN canonical variate, the six *APOE* genotypes (ɛ2/2, ɛ2/3, ɛ3/3, ɛ2/4, ɛ3/4, and ɛ4/4), and six interaction terms capturing the non-linear association between each of the six *APOE* genotypes and the given HC or DN pattern, controlling for age. We thus obtained a total of 100 logistic models that sought to explain variance in family history of ADRD as a function of these 14 parameters. We performed the analogous permutation analysis (described above) to assess the robustness of each of the 14 regression coefficients derived from these 100 logistic models.

### Latent factor analysis of brain-behaviour associations

To finally distill latent factor embeddings of brain-behaviour associations from our HC-DN population signatures, we used the classical linear dimensionality reduction method principal component analysis (PCA) [173]. PCA was a natural choice of method to uncover linearly independent groupings of risk factors with similar relatedness to HC-DN co-variation patterns. Latent factors uncovered by the PCA are naturally ordered from most to least important which allows us to select candidate principles of brain-behaviour association that account for the most inter-individual variance.

We started by computing the Pearson’s correlation between the z-scored canonical variate matrix (number of participants x 50 canonical variates) and the z-scored risk factor matrix (number of participants x 63 risk factors). We obtained a new matrix *M* ∈ R^63 × 50^, which represented the Pearson’s correlation coefficients between the 63 risk factors and the 50 canonical variates. We next decomposed *M* into latent factor groupings by using singular value decomposition (SVD). Every correlation coefficient in *M* had already been z-scored to abide by zero mean and unit variance prior to computing the SVD, as per common practice [174]. More formally, solving the SVD problem took the following form:

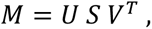

where *U* is a 63 x 63 orthonormal matrix, *S* is a 63 x 50 diagonal matrix carrying the singular values and *V* is a 50 x 50 orthonormal matrix carrying singular vectors.

We retained the top three singular vectors and expressed our correlation matrix in terms of the dot product *US* ∈ *R*^63 × 3^ to be able to represent the latent-factor projections of *M* onto the new three-dimensional latent space. Doing so, we obtained the distinct expression levels of the 63 risk factors for each of the top three brain-behaviour associations axes (i.e., principal component expressions). These three axes are by construction orthonormal and rank-ordered, representing an uncorrelated partition of the overall variance in brain-behaviour association. The leading axis captured the largest fraction of variance and was therefore the most explanatory, as reflected by its associated singular value.

We then conducted an acid test of the robustness of the PCA solution by performing a rigorous split-half reliability assessment. We derived two random subsets of equal size (N=18,645) from the original sample and re-computed the Pearson’s correlation matrix *M* for each random subset separately. SVD was then performed on both matrices in parallel according to the procedure described above. We retained the same number of top three singular vectors and expressed each correlation matrix in term of its projection onto its corresponding latent space. In doing so, we were able to compare the expression levels of each risk factor along the three main axes of brain-behaviour associations derived from each random subset. If the PCA solution is robust, similar groups of risk factors should be emphasized along corresponding dimensions which, in turn, should explain similar fractions of the total variance.

Based on the desire to audit our cohort analysis for sex-specific associations, we computed the Pearson’s correlation matrix *M* in males and females separately and repeated the PCA procedure described above for each group. Once more, we retained the top three singular vectors and expressed the correlation matrices in term of their projection onto their corresponding latent embedding. We compared the expression levels of the risk factors along corresponding latent dimensions to highlight sex-specific brain-behaviour associations. In the absence of major sex differences, similar groups of risk factors should be emphasized along analogous dimensions which should correspondingly explain similar fractions of the total variance.

---

**Extended Data Figure 1.**
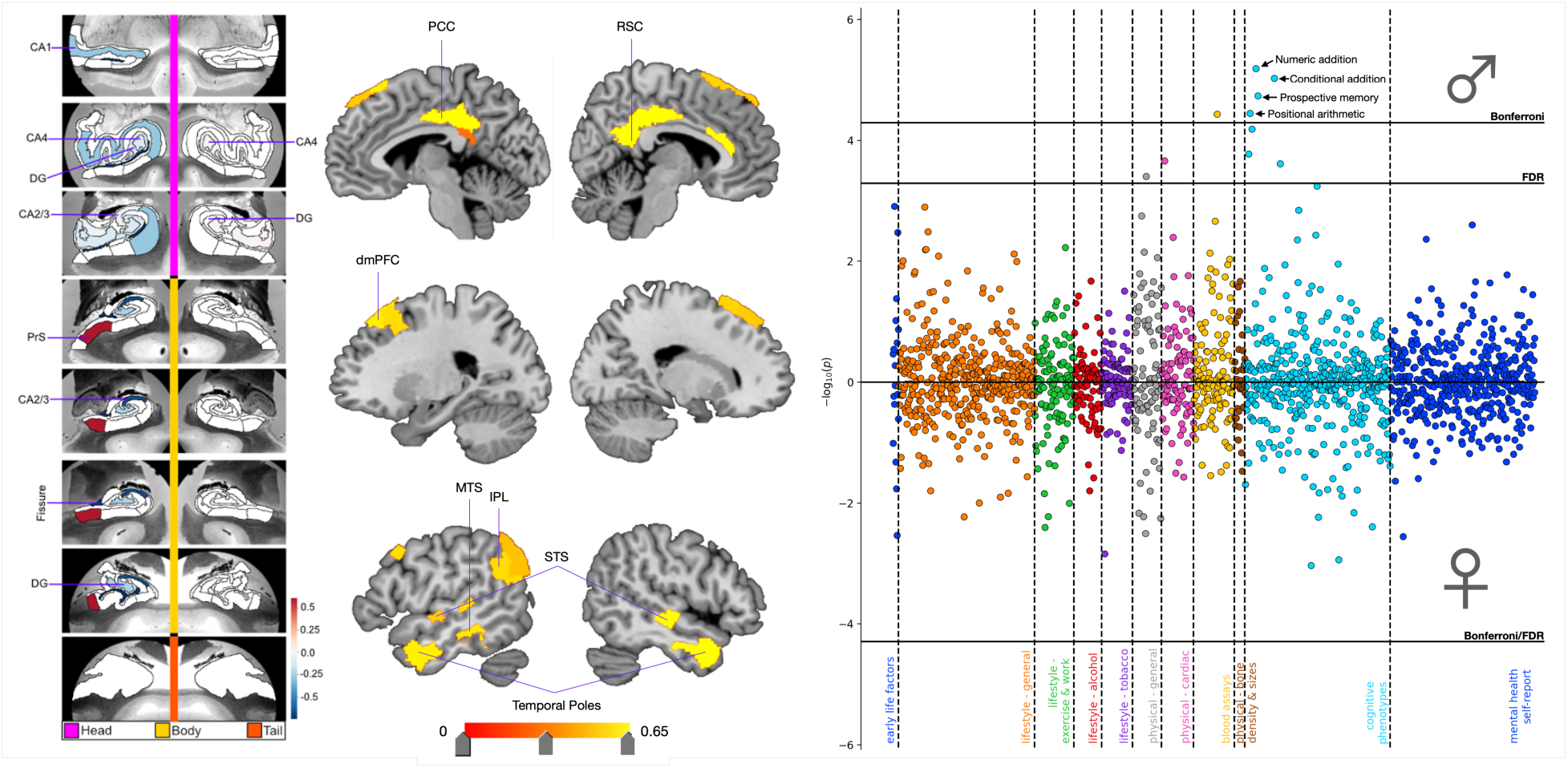
ADRD-related divergences in HC and DN subregions for mode 2 and the associated phenome-wide profile. Shown here are ADRD-related subregion divergences for mode 2 for the HC (leftmost panel) and DN (central panel). We identified 10 HC hits, most of them located in the left hemisphere. The strongest HC divergences were observed for the presubiculum, hippocampal fissure, and CA2/3. We found corresponding DN hits in posterior midline structure (posterior cingulate cortex and restrosplenial cortex), the dorsomedial prefrontal cortex, and the posterior and temporal cortices. In males and females separately, we regressed *APOE* dosage on HC and DN co-variation patterns from mode 2. We then used these sex-specific models to predict *APOE* dosage based on inter-individual expressions of mode 2. The right panel displays the Miami plot for the correlations between *APOE* scores in the context of mode 2 and the portfolio of UKB phenotypes for males (upper half) and females (lower half). We found significant associations with sub- questions from the fluid intelligence battery that were unique to males. CA = cornu amonis, DG = granule cell layer of the dentate gyrus, PrS= presubiculum, PCC = posterior cingulate cortex, RSC = retrosplenial cortex, dmPFC = dorsomedial prefrontal cortex, IPL = inferior parietal lobule, MTS = middle temporal sulcus, and STS = superior temporal sulcus, FDR = false discovery rate correction.

**Extended Data Figure 2.**
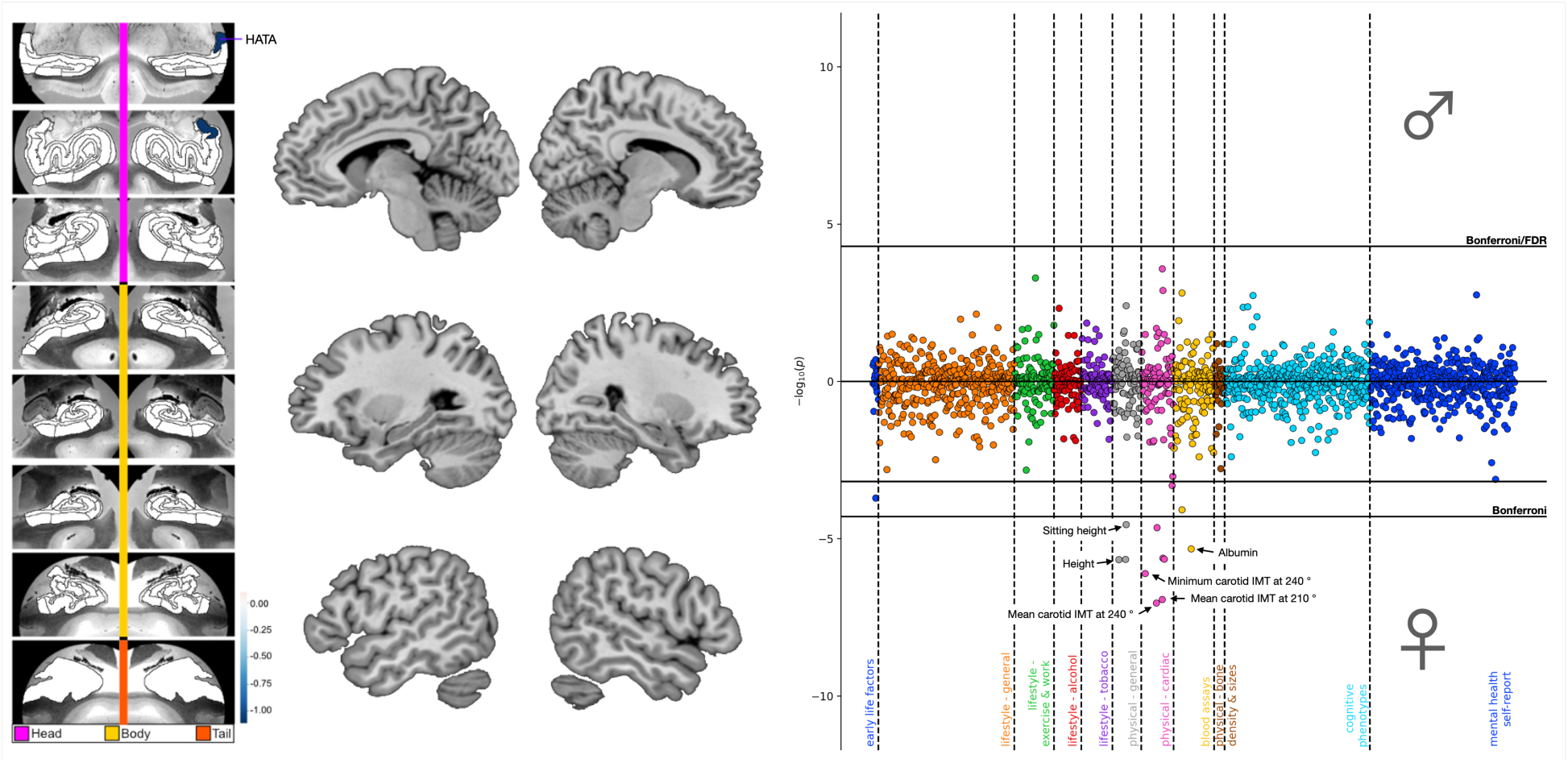
ADRD-related divergences in HC and DN subregions for mode 6 and the associated phenome-wide profile. Shown here are ADRD-related subregion divergences for mode 6 for the HC (leftmost panel) and DN (central panel). We identified 1 HC hit to the hippocampus-amygdala transition area with no concurrent DN divergences. In males and females separately, we regressed *APOE* dosage on HC and DN co-variation patterns from mode 6. We then used these sex-specific models to predict *APOE* dosage based on inter-individual expressions of mode 6. The right panel displays the Miami plot for the correlations between *APOE* scores in the context of mode 6 and the portfolio of UKB phenotypes for males (upper half) and females (lower half). We found significant associations with physical phenotypes and blood assays that were unique to females. HATA = hippocampus-amygdala transition area, IMT = intima-medial thickness, FDR = false discovery rate correction.

**Extended Data Figure 3.**
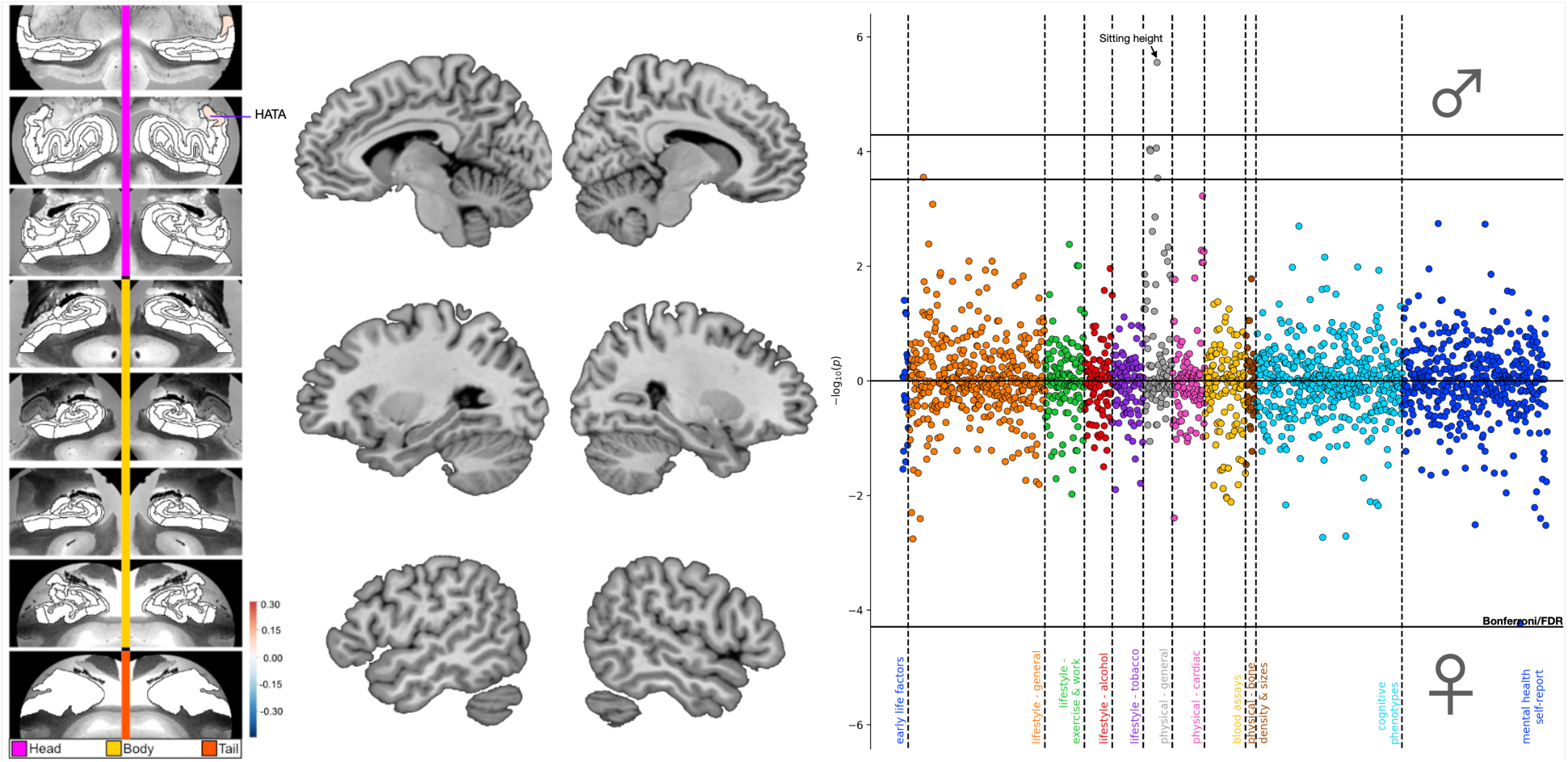
ADRD-related divergences in HC and DN subregions for mode 10 and the associated phenome-wide profile. Shown here are ADRD-related subregion divergences for mode 10 for the HC (leftmost panel) and DN (central panel). We identified 1 HC hit to the hippocampus-amygdala transition area with no concurrent DN divergences. In males and females separately, we regressed *APOE* dosage on HC and DN co-variation patterns from mode 10. We then used these sex-specific models to predict *APOE* dosage based on inter-individual expressions of mode 10. The right panel displays the Miami plot for the correlations between *APOE* scores in the context of mode 10 and the portfolio of UKB phenotypes for males (upper half) and females (lower half). We found one significant association with sitting height unique to males. HATA = hippocampus-amygdala transition area, FDR = false discovery rate correction.

**Extended Data Figure 4.**
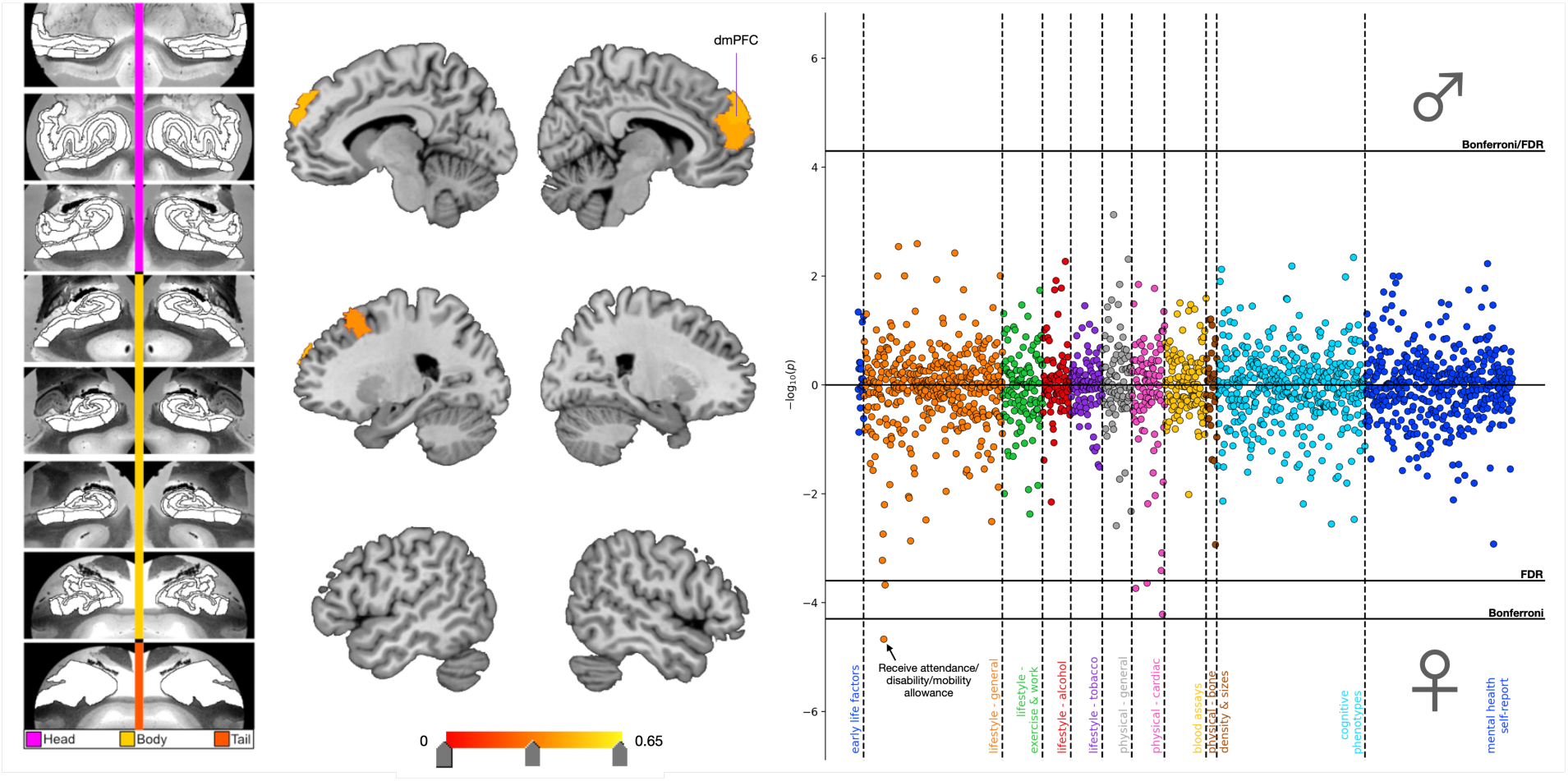
ADRD-related divergences in HC and DN subregions for mode 4 and the associated phenome-wide profile. Shown here are ADRD-related subregion divergences for mode 4 for the HC (leftmost panel) and DN (central panel). We identified 4 DN hits to the dorsomedial prefrontal cortex with no concurrent HC divergences. In males and females separately, we regressed *APOE* dosage on HC and DN co-variation patterns from mode 4. We then used these sex-specific models to predict *APOE* dosage based on inter-individual expressions of mode 4. The right panel displays the Miami plot for the correlations between *APOE* scores in the context of mode 4 and the portfolio of UKB phenotypes for males (upper half) and females (lower half). We found one significant association with receiving an attendance, disability, or mobility allowance that was unique to females. dmPFC = dorsomedial prefrontal cortex, FDR = false discovery rate correction.

**Extended Data Figure 5.**
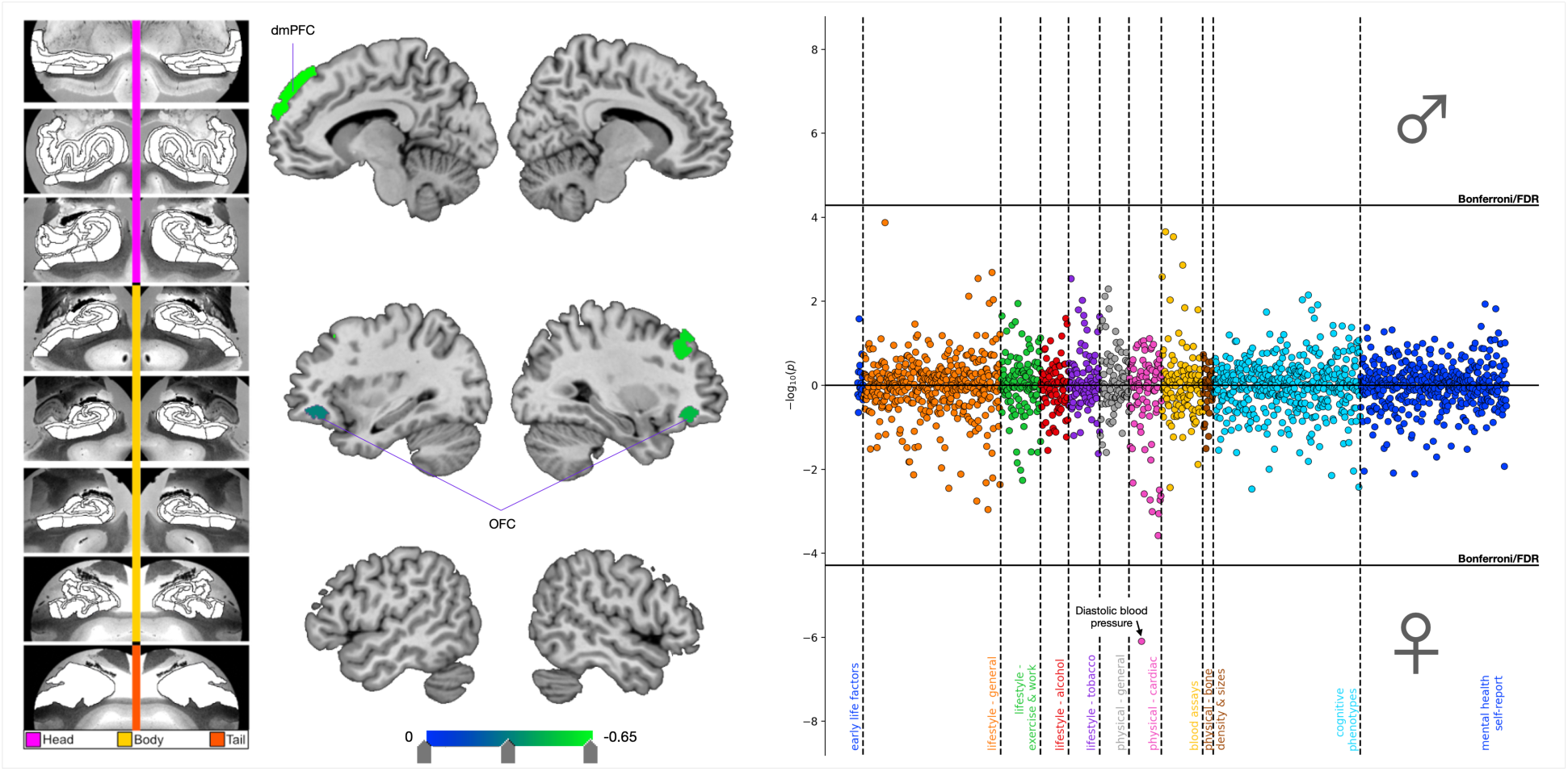
ADRD-related divergences in HC and DN subregions for mode 7 and the associated phenome-wide profile. Shown here are ADRD-related subregion divergences for mode 7 for the HC (leftmost panel) and DN (central panel). We identified 9 DN hits to the frontal lobe with no concurrent HC divergences. In males and females separately, we regressed *APOE* dosage on HC and DN co-variation patterns from mode 7. We then used these sex-specific models to predict *APOE* dosage based on inter-individual expressions of mode 7. The right panel displays the Miami plot for the correlations between *APOE* scores in the context of mode 7 and the portfolio of UKB phenotypes for males (upper half) and females (lower half). We found one significant association with diastolic blood pressure that was unique to females. dmPFC = dorsomedial prefrontal cortex, OFC = orbitofrontal cortex, FDR = false discovery rate correction.

**Extended Data Figure 6.**
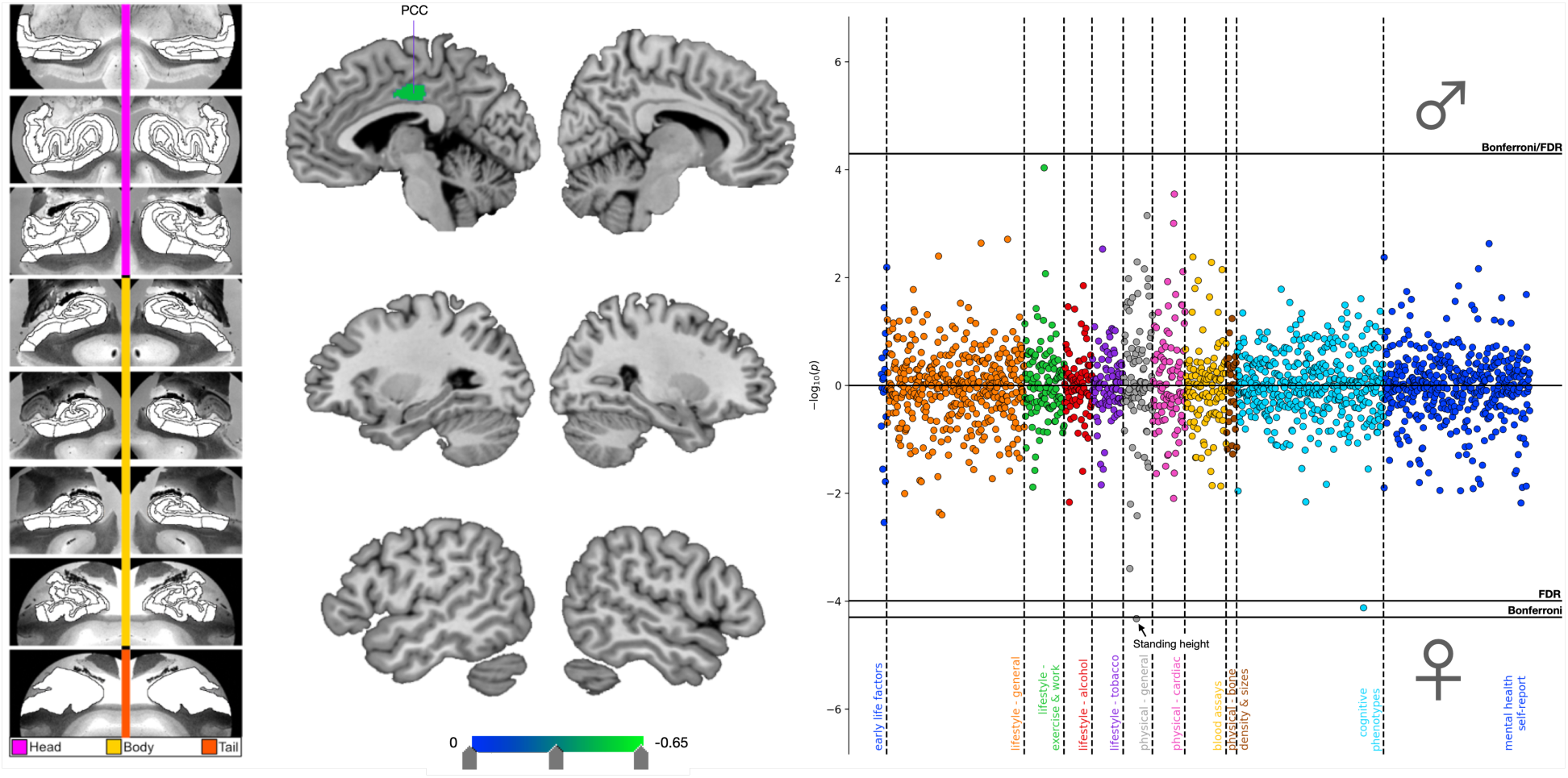
ADRD-related divergences in HC and DN subregions for mode 11 and the associated phenome-wide profile. Shown here are ADRD-related subregion divergences for mode 11 for the HC (leftmost panel) and DN (central panel). We identified 1 DN hit to posterior cingulate cortex with no concurrent HC divergences. In males and females separately, we regressed *APOE* dosage on HC and DN co- variation patterns from mode 11. We then used these sex-specific models to predict *APOE* dosage based on inter-individual expressions of mode 11. The right panel displays the Miami plot for the correlations between *APOE* scores in the context of mode 11 and the portfolio of UKB phenotypes for males (upper half) and females (lower half). We found one significant association with standing height that was unique to females. PCC = posterior cingulate cortex, FDR = false discovery rate correction.

**Extended Data Figure 7.**
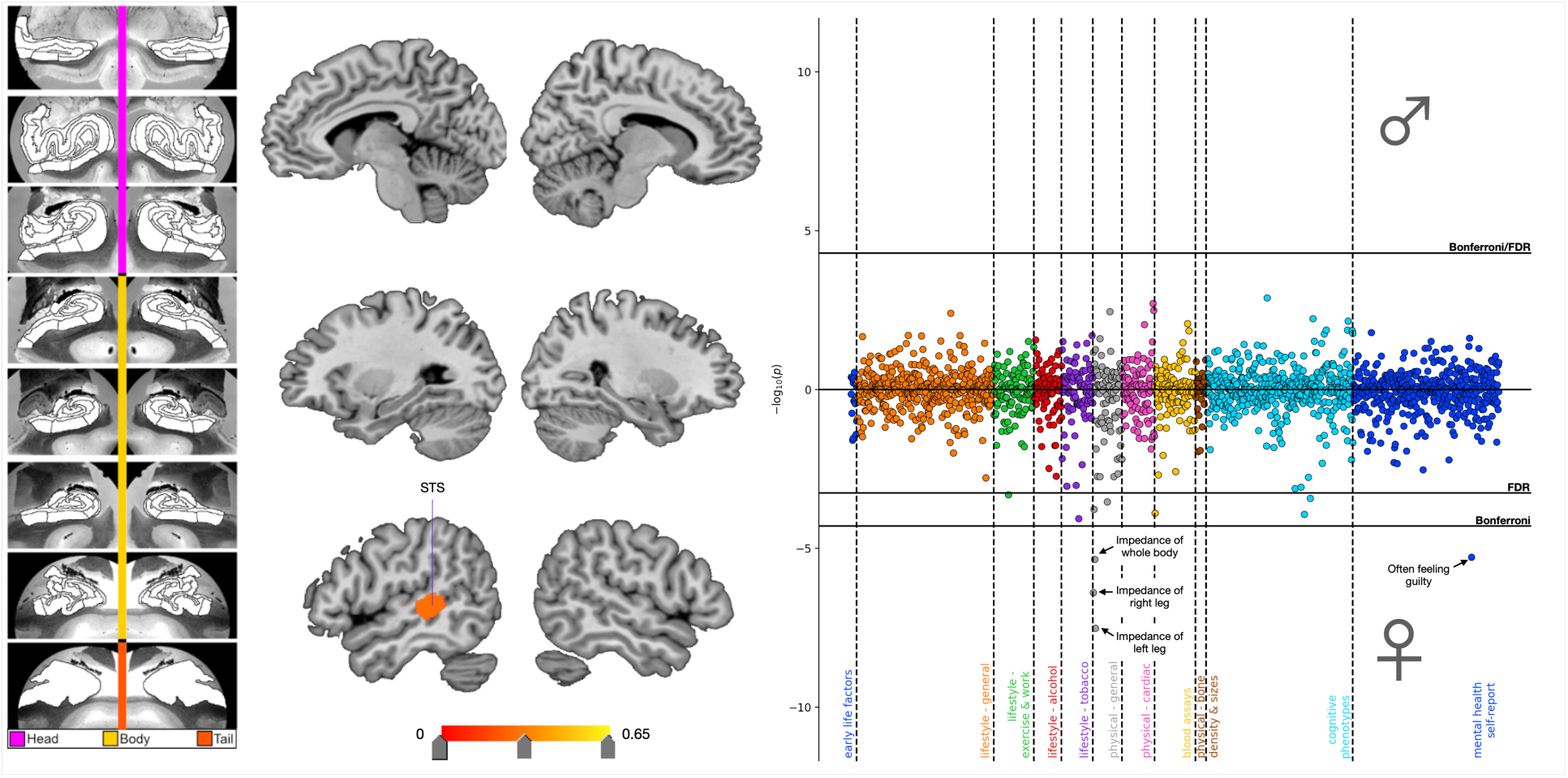
ADRD-related divergences in HC and DN subregions for mode 13 and the associated phenome-wide profile. Shown here are ADRD-related subregion divergences for mode 13 for the HC (leftmost panel) and DN (central panel). We identified 1 DN hit to the superior temporal sulcus with no concurrent HC divergences. In males and females separately, we regressed *APOE* dosage on HC and DN co- variation patterns from mode 13. We then used these sex-specific models to predict *APOE* dosage based on inter-individual expressions of mode 13. The right panel displays the Miami plot for the correlations between *APOE* scores in the context of mode 13 and the portfolio of UKB phenotypes for males (upper half) and females (lower half). We found significant associations with physical measurements related to height as well as feelings of guilt that were unique to females. STS = superior temporal sulcus, FDR = false discovery rate correction.

**Extended Data Figure 8.**
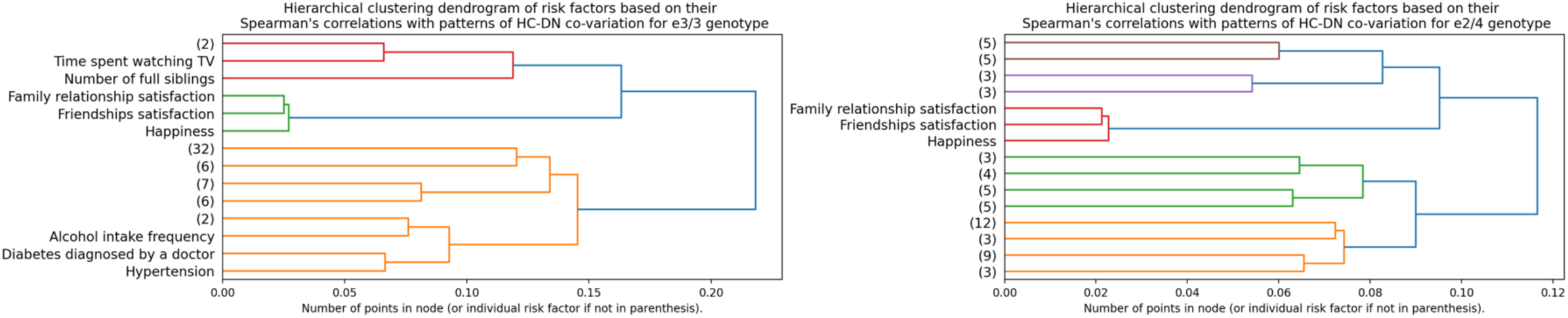
The interaction of *APOE* ɛ3/3 and ɛ2/4 with patterns of HC-DN co- variation is related to happiness and satisfaction with friendships and family. We multiplied the population-wide HC and DN co-variation patterns by *APOE* genotypes ɛ3/3 (N=22,129) and ɛ2/4 (N=885) such that participants who do not carry a given genotype were zeroed out. We then computed the Spearman’s correlations between these two new vectors and the 63 pre-selected Alzheimer’s disease risk factors to test for risk-anatomy links. We performed an agglomerative clustering analysis on these Spearman’s correlations, which consists in repeatedly merging Spearman’s correlations with similar variance together until all observations are merged into a single cluster. Here are shown the dendrograms which indicate the distance between each cluster identified when retaining three levels of branching for *APOE* ɛ3/3 (leftmost panel) and ɛ2/4 (rightmost panel). In the clustering analysis for both ɛ3/3 and ɛ2/4, we observed the emergence of the social cluster that comprised ‘family relationship satisfaction’, ‘happiness’, and ‘friendships satisfaction’ as what the case for most other *APOE* gene variants. In ɛ3 homozygotes, we also observed the emergence of a cardiovascular cluster that comprised ‘alcohol intake frequency’, ‘diabetes diagnosed by a doctor’, and ‘hypertension’. We thus showed that social phenotypes formed important risk-anatomy links in ɛ3 homozygotes and ɛ2/4 carriers.

**Extended Data Figure 9.**
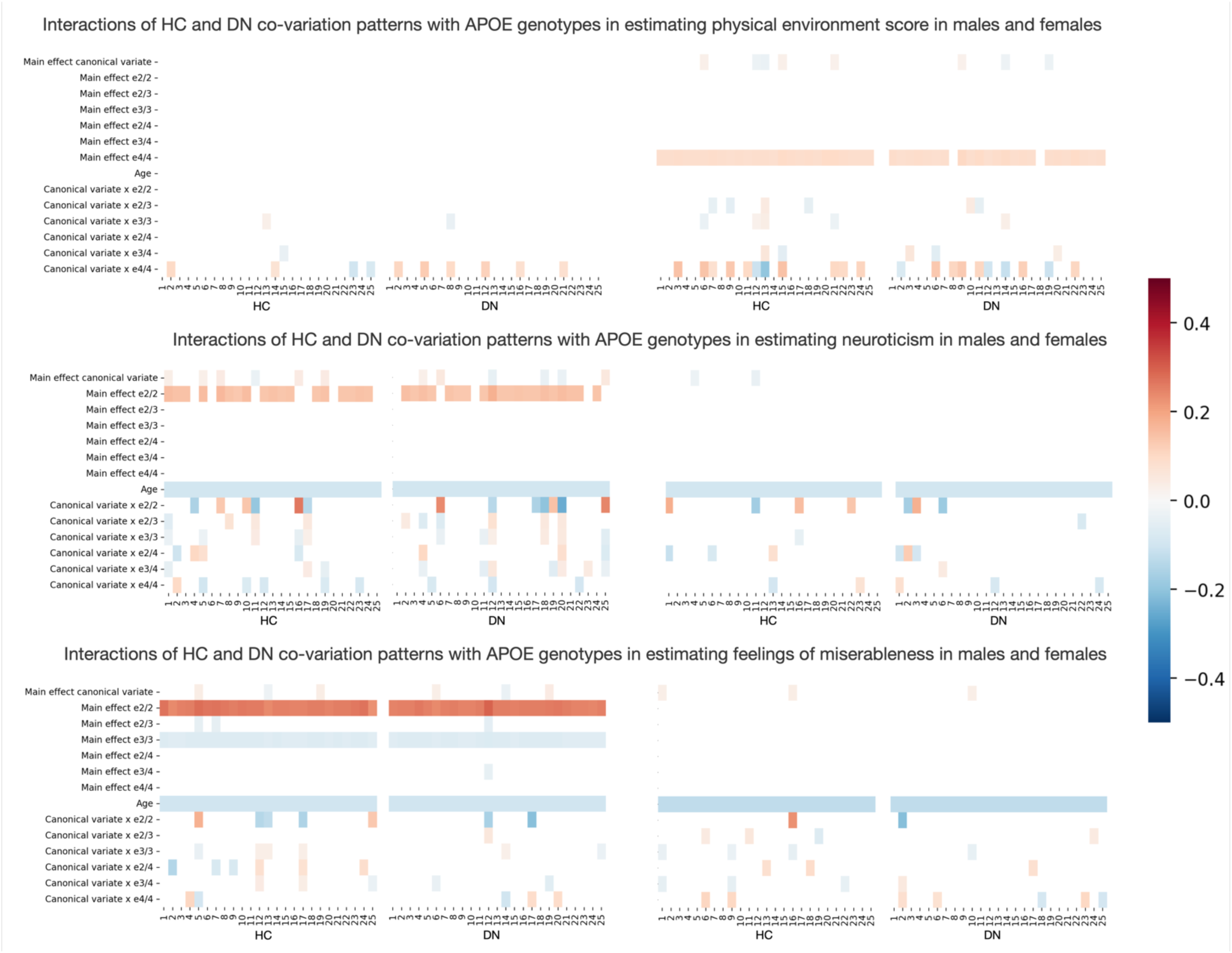
Brain-*APOE* ɛ4/4 interaction explains variance in physical environment score while ɛ2/2 is linked to neuroticism and feelings of miserableness in males. We tested whether HC-DN signatures interacted with *APOE* genotypes in explaining variance on the 63 pre-selected ADRD risk factors. Each risk factor was individually regressed on a single HC or DN pattern from a given mode of HC-DN co-variation, resulting in 50 different linear models per risk factor. Each model took as parameters the main effect of a given HC or DN pattern, the main effects of the six *APOE* genotypes (i.e., ɛ2/2, ɛ2/3, ɛ3/3, ɛ2/4, ɛ3/4, and ɛ4/4), and the interaction between each of the six *APOE* genotype and the given HC or DN pattern, controlling for age. Separate analyses were run for males (leftmost plots) and females (rightmost plots). Each column on the heat maps thus represents the coefficients for a single linear regression model. The first 25 columns show the coefficients for HC patterns, whereas the last 25 columns show the coefficients for DN patterns. We assessed the robustness of our findings by comparing each coefficient to empirically built null distributions obtained through permutation testing. Only the coefficients that were statistically different from their respective null distributions in 95% of the time are presented. We displayed the modifiable risk factors for which the strongest brain-APOE interactions were observed. On the upper panels, we show that *APOE* ɛ4/4 preferentially interacts with HC and DN co-variations patterns in estimating physical environment score, a measure of environmental factors that have an impact on the quality of life in an area. We also found a significant main effect of ɛ4/4 unique to females. On the middle panels, we show that *APOE* ɛ2/2 interacts with HC and DN canonical variates in estimating neuroticism, with an accompanying significant main effect of ɛ2/2 found in males. Similar, on the lower panels, we show a strong main effect of *APOE* ɛ2/2 in estimating feelings of miserableness, a personality-trait correlated with neuroticism, in males. These interactions profiles can be taken to suggest that ɛ4 homozygotes are biologically susceptible to environmental factors that may impact quality of life. We also reiterated the link between neuroticism and ɛ2 by showing a sparse pattern of brain-gene interactions unique to ɛ2 homozygotes, with stronger main effects found in males.

**Extended Data Figure 10:**
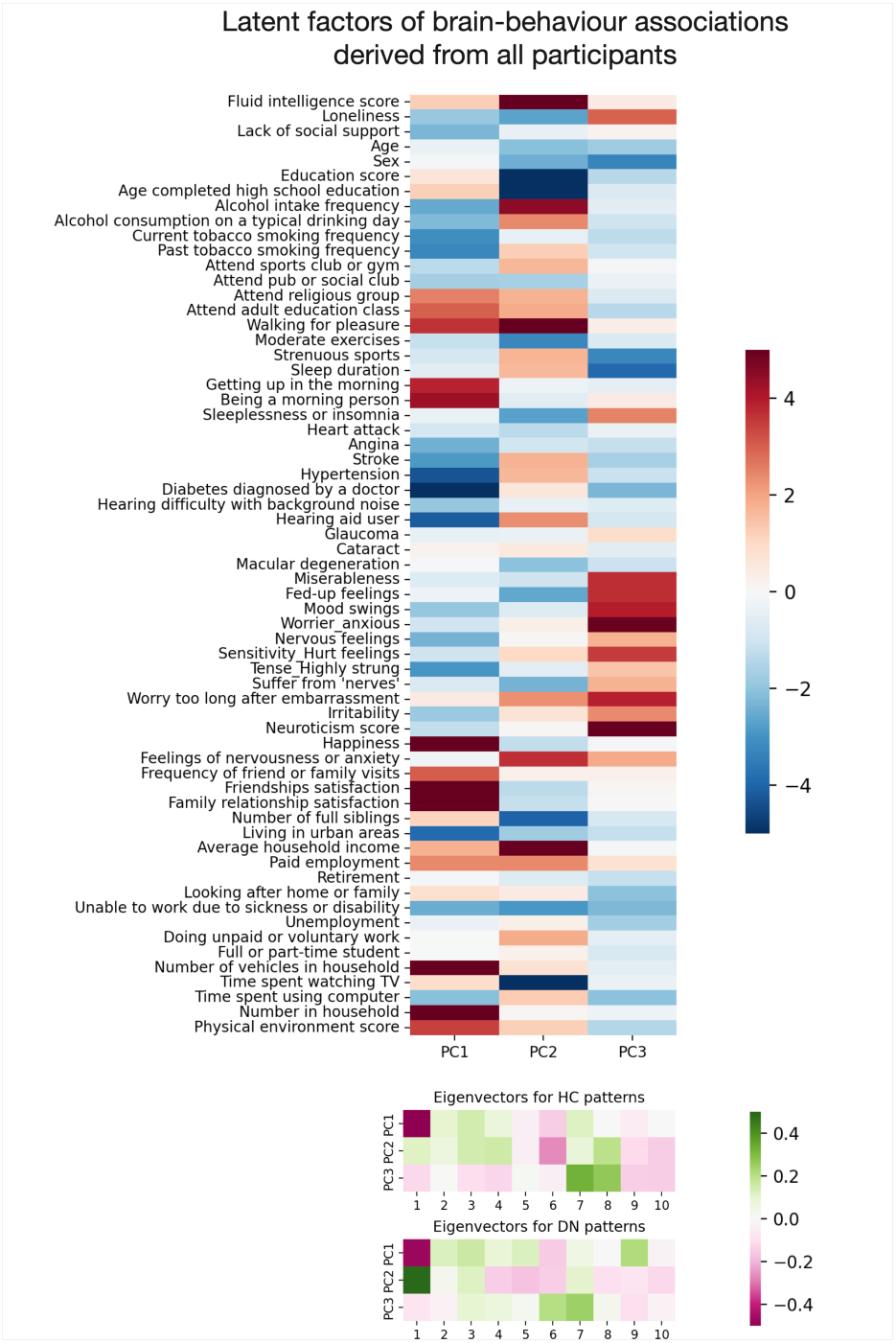
Latent factors of brain-behaviour associations emphasize satisfaction with social relationships, socioeconomic status, and neuroticism-related traits. We conducted an exploratory principal component analysis (PCA) to disentangle latent factor of brain-behaviour association in our UK Biobank sample. We first computed the Pearson’s correlations between the 25 pairs of co-variation patterns from the HC and DN sides and the 63 pre-selected ADRD risk factors. We then ran singular value decomposition on the risk by canonical variates matrix (X63 x 50) and retained the 3 first principal components (PCs) that explained ∼16.3%, ∼12.8%, and ∼8.2% of the total variance in the data, respectively. The upper plot displays the projections of the Pearson’s correlations onto each of the three main axes of brain-behaviour associations. The lower plot displays the eigenvectors for the top ten HC and DN co-variation patterns. The first axis of brain-behaviour associations emphasizes phenotypes from the social cluster previously identified on the clustering analysis of risk-anatomy links (Fig. 4), i.e., ‘family relationship satisfaction’, ‘happiness’, and ‘friendships satisfaction’. The second axis rather accented socioeconomic risk factors such as fluid intelligence, education, alcohol intake, average household income and time spent watching television. Lastly, the third axis of brain-behaviour associations separates neuroticism-related items (e.g., ‘miserableness’, ‘fed-up feelings’, ‘mood-swings’, and ’worrier/anxious’) from the rest of the risk factors.

