## Supplementary Figures 1 & 2 for "*APOE* ɛ2 vs *APOE* ɛ4 dosage shows sex-specific links to hippocampus-default network subregion co-variation"

### Supplementary information

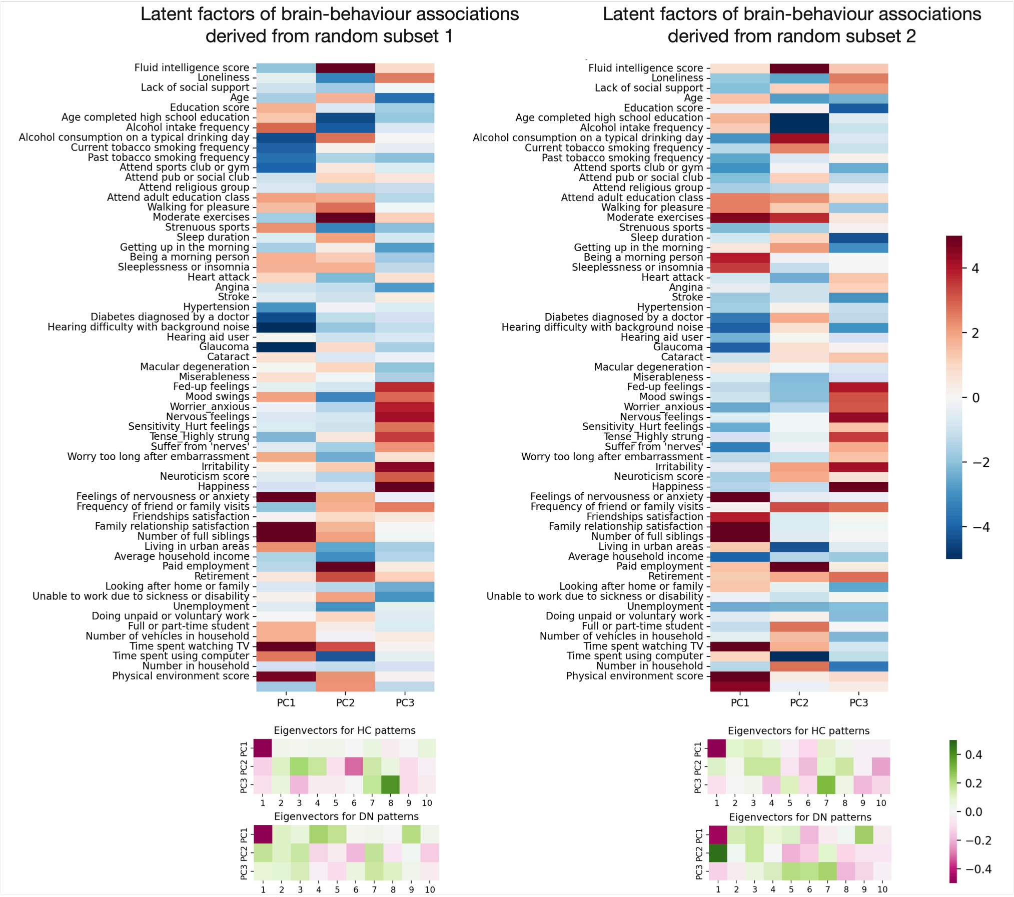


**Supplementary Figure 1: Split-half reliability assessment of the principal component solution**

We assessed the robustness of the derived brain-behaviour associations axes by repeating the principal component procedure in two random subsets of equal size (N=18,645) derived from the original data. For each subset, we first computed the Pearson's correlations between the 25 pairs of co-variation patterns from the HC and DN sides and the 63 pre-selected ADRD risk factors. We then ran singular value decomposition on the risk by canonical variates matrix ($X_{63 \times 50}$) and retained the 3 first principal components (PCs). The PCs obtained from the first random subset had explained variance of ~13.9%, 11.1%, and 7.9%, respectively. The PCs obtained from the second random subset had explained variance of ~15.3%, 11.2%, and 9.1%, respectively. The upper plots display the projections of the Pearson's correlations onto each of the three axes of brain-behaviour associations for the two random subsets. The lower plots display the eigenvectors for the top ten HC and DN co-variation patterns. Looking at the projections of the Person's correlation onto the principled axis of brain-behaviour associations, we can observe a good reliability between the two subsets with the same sets of risk factors scoring the strongest along corresponding axes.

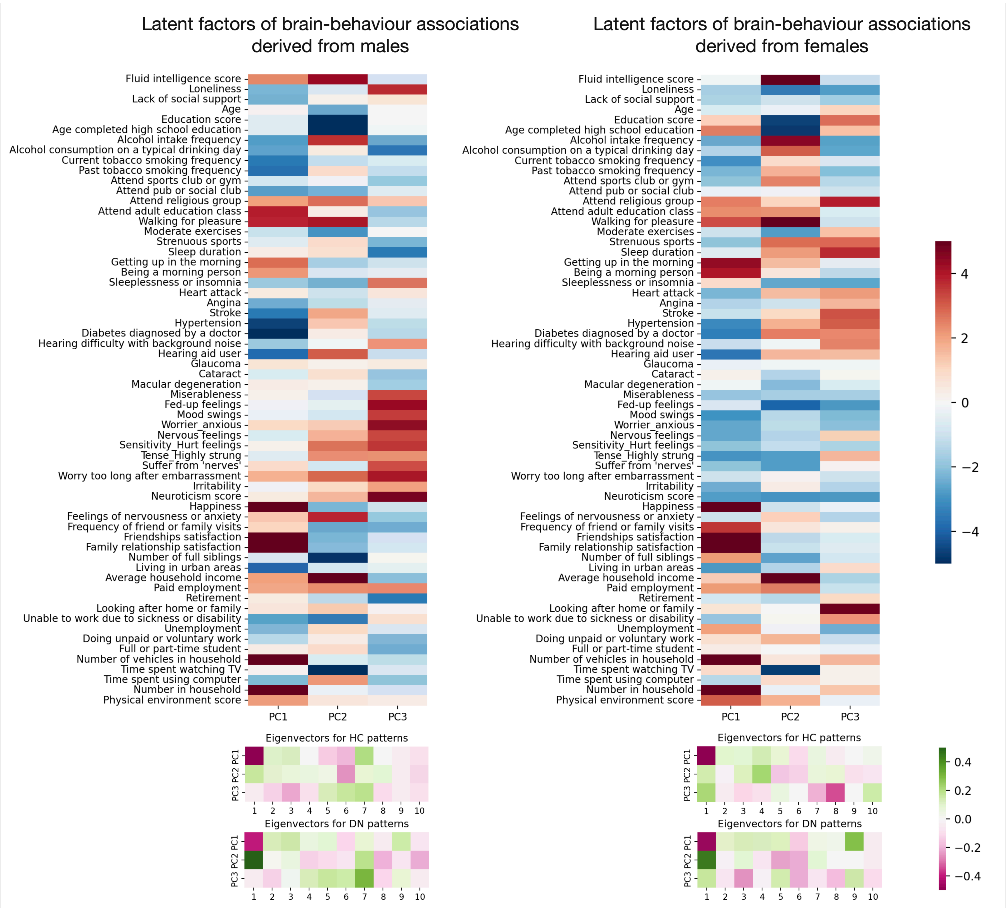


**Supplementary Figure 2: Neuroticism-related items expressed distinctive brain-behaviour associations in males and females**

We repeated the principal component analysis in males (left; N=17,561) and females right; N=19,730) separately. In each sex, we first computed the Pearson's correlations between the 25 pairs of co-variation patterns from the HC and DN sides and the 63 pre-selected ADRD risk factors. We then ran singular value decomposition on the risk by canonical variates matrix ($X_{63 \times 50}$) and retained the 3 first principal components (PCs). The PCs obtained from males had explained variance of ~14.6%, ~11.9%, and ~9.6%, respectively. The PCs obtained from females had explained variance of ~14.6%, ~11.9%, and ~7.4%, respectively. The upper plots display the projections of the Pearson's correlations onto each of the three axes of brain-behaviour associations for the two sexes. The lower plots display the eigenvectors for the top ten HC and DN co-variation patterns. The projections of the Pearson's correlations onto the two first axes of brain-behaviour association were roughly the same in males and females. In contrast, neuroticism-related items were only emphasized on the third axis of brain-behaviour association in males. We thus supplemented our population analysis by showing that the relationship between neuroticism and patterns of HC-DN co-variation was mainly male-specific.
